# NUDT5 regulates purine metabolism and thiopurine sensitivity by interacting with PPAT

**DOI:** 10.1101/2025.03.29.646096

**Authors:** Zheng Wu, Phong T Nguyen, Varun Sondhi, Run-Wen Yao, Tao Dai, Jui-Chung Chiang, Zengfu Shang, Feng Cai, Ling Cai, Jing Zhang, Mya D Moore, Islam Alshamleh, Xiangyi Li, Tamaratare Ogu, Lauren G Zacharias, Rainah Winston, Joao S Patricio, Xandria Johnson, Wei-Min Chen, Qian Cong, Thomas P Mathews, Yuanyuan Zhang, Ralph J DeBerardinis

## Abstract

Cells generate purine nucleotides through both de novo purine biosynthesis (DNPB) and purine salvage. Purine accumulation represses energetically costly DNPB through feedback inhibition of the enzymatic steps that produce the precursor phosphoribosylamine. Excessive DNPB is associated with human diseases including neurological dysfunction and hyperuricemia. However, the mechanisms explaining how cells balance DNPB and purine salvage are incompletely understood. Data from a genome-wide CRISPR loss-of-function screen and extensive stable isotope tracing identified Nudix hydrolase 5 (NUDT5) as a suppressor of DNPB during purine salvage. NUDT5 ablation allows DNPB to persist in the presence of either native purines or thiopurine drugs; this renders NUDT5-deficient cells insensitive to thiopurine treatment. Surprisingly, this regulation occurs independently of NUDT5’s known function in hydrolyzing ADP-ribose to AMP and ribose-5-phosphate. Rather, NUDT5 interacts with phosphoribosyl pyrophosphate amidotransferase (PPAT), the rate-limiting enzyme in DNPB that generates phosphoribosylamine. Upon induction of purine salvage, the PPAT-NUDT5 interaction is required to trigger disassembly of the purinosome, a cytosolic metabolon involved in efficient DNPB. Mutations that disrupt NUDT5’s interaction with PPAT but leave its catalytic activity intact permit excessive DNPB during purine salvage, inducing thiopurine resistance. Collectively, our findings identify NUDT5 as a regulator governing the balance between DNPB and purine salvage, underscoring its impact on nucleotide metabolism and efficacy of thiopurine treatment.

## Introduction

Cells use both de novo purine biosynthesis (DNPB) and salvage to maintain purine nucleotide pools(*1, 2*). These two pathways are coordinately regulated to ensure purine homeostasis. Inhibiting DNPB promotes purine salvage as a compensatory mechanism(*3*), while activating purine salvage represses DNPB to prevent excessive synthesis(*4*). Perturbation of purine homeostasis can disturb the purine-pyrimidine balance, consequently leading to replication stress, DNA damage, and cell cycle arrest(*5*). Mutations causing hyperactivity of phosphoribosylpyrophosphate synthetase (PRPS1), the enzyme that produces phosphoribosyl pyrophosphate (PRPP) at the initial step of purine synthesis, drive excessive DNPB through failed feedback inhibition(*6*). Patients with these pathogenic *PRPS1* variants present with hyperuricemia/gout and neurological dysfunction, underscoring the physiological importance of governing purine nucleotide synthesis(*7*).

The ability to engage either DNPB or purine salvage is pivotal for cellular adaptation to stress, including energetic stress. In contrast to purine salvage, DNPB is energetically demanding and requires metabolic input from several different pathways, including the pentose phosphate pathway, nonessential amino acid metabolism, the folate cycle and oxidative mitochondrial metabolism(*1*). We previously demonstrated that impairment of mitochondrial respiration by mutation or electron transport chain (ETC) inhibitors drives a shift from DNPB to purine salvage, preventing unnecessary energy expenditure and enabling cancer cell growth in culture and in vivo(*8*). However, a complete understanding of the mechanisms mediating the interplay between DNPB and purine salvage is lacking.

Substantial evidence, including mass spectrometry imaging at single-cell resolution, indicates that DNPB enzymes associate into a cytosolic multiprotein complex called the purinosome. This complex is thought to facilitate rapid purine synthesis by channeling intermediates from one enzyme to the next, limiting their diffusion away from the complex(*9–11*). Purinosome formation is dynamically regulated by the availability of salvageable purines. Purine deprivation induces purinosome formation to enhance DNPB, and repletion of purine bases or nucleosides leads to rapid purinosome dispersal(*9*). Thus, purinosome disassembly is part of the mechanism by which purine salvage inhibits DNPB. While several regulators of purinosome assembly have been identified (*1, 12–14*), the mechanisms underlying purinosome disassembly in response to induction of purine salvage are not well understood.

Our search for genes involved in regulating the balance between purine metabolic pathways identified *NUDT5*, which encodes the ADP-ribose (ADPR) hydrolase Nudix Hydrolase 5, as a negative regulator of DNPB during purine salvage. Surprisingly, NUDT5’s catalytic activity was dispensable for this effect. Rather, we observed that NUDT5 associates with phosphoribosyl pyrophosphate amidotransferase (PPAT), the rate-limiting enzyme in DNPB, and accelerates purinosome disassembly during purine salvage(*1, 15*).

## Results

### Ablation of NUDT5 confers cellular resistance to thiopurines

To identify regulators of purine salvage, we mined the data from a previous whole-genome CRISPR screen using 6-thioguanine (6-TG)(*16*). While 6-TG is not directly toxic to cells, it can be converted to thioguanine nucleotides such as 6-thio-GMP (6-TGMP) by the purine salvage enzyme hypoxanthine-guanine phosphoribosyltransferase-1 (HPRT1). These thioguanine nucleotides are incorporated into the genome during DNA synthesis, causing DNA damage and cell death (Fig. 1A). Therefore, 6-TG-induced toxicity depends on HPRT1 enzymatic activity. As expected, gRNAs targeting *HPRT1* are positively selected in 6-TG-treated cells (fig. S1A). Interestingly, gRNAs targeting Nudix Hydrolase 5 (*NUDT5*) were positively selected nearly as much as gRNAs targeting *HPRT1* (fig. S1A). NUDT5 catalyzes the conversion of ADP-ribose (ADPR) to AMP and ribose-5-phosphate (R5P)(*17*). Because R5P is a precursor of phosphoribosyl pyrophosphate (PRPP), which provides the activated ribose for purine synthesis by HPRT1 (Fig. 1A), NUDT5 has a plausible role in 6-TG toxicity. Therefore, to validate the screening result, we used CRISPR-Cas9 technologies to delete *HPRT1* or *NUDT5* in HeLa cervical carcinomas cells and A549 lung adenocarcinomas cells (fig. S1B). Loss of NUDT5 or HPRT1 increased the level of their substrates ADPR and hypoxanthine, respectively (fig. S1C,D). Consistent with the screen results, deficiency of either HPRT1 or NUDT5 conferred resistance to 6-TG and the related thiopurine 6-mercaptopurine (6-MP) (Fig. 1B,C and fig. S1E,F).

**Figure 1.**
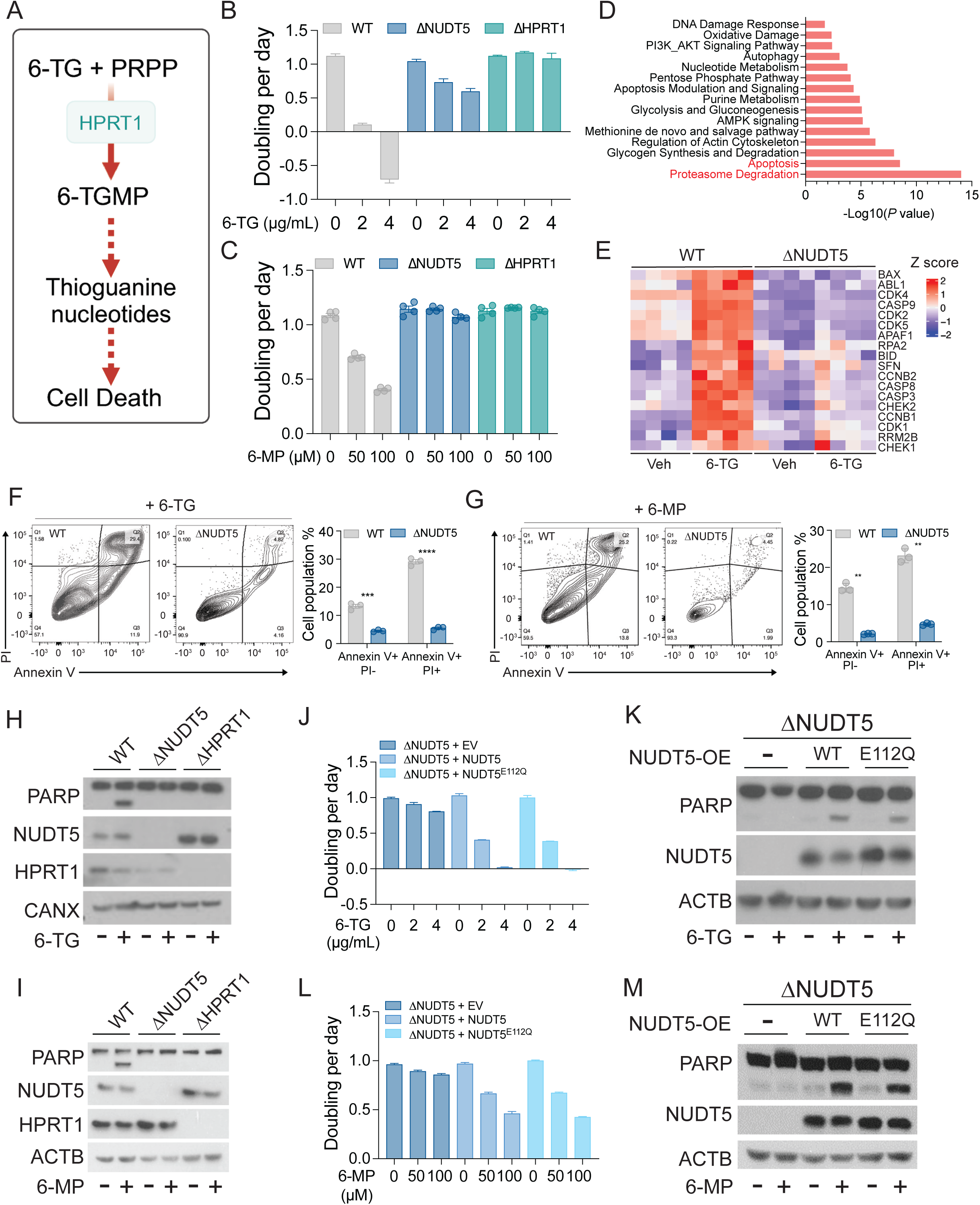
NUDT5 regulates cellular sensitivity to thiopurines independently of its catalytic function. **A.** Simplified schematic illustrating 6-TG metabolism-mediated cell death. **B-C.** Growth rates of WT, ΔNUDT5, and ΔHPRT1 HeLa cells treated with the indicated doses of 6-TG (**B**) or 6-MP (**C**). Data are from one of three independent experiments. **D.** Proteomics analysis showing upregulated pathways in 6-TG-treated WT cells compared to vehicle-treated WT, vehicle-treated ΔNUDT5 and 6-TG-treated ΔNUDT5 cells. **E.** Heatmap showing abundance of proteins in the apoptosis pathway. **F-G.** Apoptosis analysis in WT and ΔNUDT5 HeLa cells treated with 0.5 µg/mL 6-TG (**F**) or 20 µM 6-MP (**G**) for 48 hours. To the left are representative contour plots. To the right are bar graphs showing the percentage of apoptotic (Annexin V+, PI −) and dead (Annexin V+, PI+) cells. Data are means from three replicates. **H-I.** Western blot assessing cleaved PARP in WT, ΔNUDT5, and ΔHPRT1 HeLa cells treated with 0.5 µg/mL 6-TG (**H**) or 20 µM 6-MP (**I**) for 24 hours. Calnexin (CANX) is the loading control. **J.** Growth rates of ΔNUDT5 HeLa cells that express empty vector (EV), WT NUDT5, or NUDT5^E112Q^ treated with the indicated doses of 6-TG. **K.** Western blot assessing cleaved PARP in ΔNUDT5 HeLa cells that express empty vector (EV), WT NUDT5, or NUDT5^E112Q^ treated with 0.5 µg/mL 6-TG for 24 hours. β-actin (ACTB) is the loading control. **L.** Growth rates of ΔNUDT5 HeLa cells that express empty vector (EV), WT NUDT5, or NUDT5^E112Q^ treated with the indicated doses of 6-MP. **M.** Western blot assessing cleaved PARP in ΔNUDT5 HeLa cells that express empty vector (EV), WT NUDT5, or NUDT5^E112Q^ treated with 20 µM 6-MP for 24 hours. β-actin (ACTB) is the loading control. Multiple t test was used for statistical analyses (**F** and **G**) ****: P < 0.0001; ***: P < 0.001; **: P < 0.01. Error bars denote SEM. BioRender was used to generate the illustration in **A**.

To broadly assess how cells respond to 6-TG in the presence and absence of NUDT5, we performed quantitative proteomics on wild-type (WT) and NUDT5-null cells treated with vehicle or a low dose of 6-TG for 24 hours. Pathway analysis revealed pronounced apoptotic and proteasome degradation signatures in WT but not NUDT5-null cells (Fig. 1D,E). Indeed, Annexin V and propidium iodide (PI) staining confirmed a large increase in apoptosis after treatment with 6-TG or 6-MP in WT relative to NUDT5-null cells (Fig. 1F,G). Thiopurines also induced cleavage of Poly(ADP-ribose) polymerase (PARP) in WT cells, indicating an apoptotic response to the drugs. This induction was abrogated by deletion of either NUDT5 or HPRT1 (Fig 1H,I). Conversely, overexpression of NUDT5 resulted in excess cleaved PARP following 6-TG treatment, underscoring the role of NUDT5 in regulating thiopurine sensitivity (fig. S1G)

### NUDT5 regulates cellular responses to thiopurines independently of its catalytic function

To test the hypothesis that NUDT5’s role in thiopurine sensitivity is to provide R5P and PRPP for purine salvage by HPRT1, we compared the effect of rescuing NUDT5-null cells with WT NUDT5 or a NUDT5 mutant (NUDT5^E112Q^) reported to lack catalytic activity(*18, 19*). As expected, ectopically expressing WT NUDT5 in NUDT5-null cells diminished ADPR levels, but expression of either an empty vector or NUDT5^E112Q^ did not, confirming the catalytic defect of NUDT5 ^E112Q^ (fig. S1H). Surprisingly, however, both WT NUDT5 and NUDT5^E112Q^ restored sensitivity to thiopurines (Fig. 1J to M). We conclude that NUDT5 regulates cellular sensitivity independently of its known catalytic function.

### Thiopurines suppress mitochondrial oxidative metabolism in a NUDT5-dependent manner

Intriguingly, our proteomics data revealed that several down-regulated pathways in 6-TG-treated WT cells involve oxidative mitochondrial metabolism (fig. S2A). Many mitochondrial electron transport chain (ETC) complex subunits and tricarboxylic acid (TCA) cycle enzymes were depleted in WT but not NUDT5-null cells following 6-TG treatment (fig. S2B), in corroboration with previous findings in acute lymphoblastic leukemia cells (ALL)(*20*). Consequently, WT cells treated with 6-TG displayed reduced oxygen consumption and reduced contributions of oxidative metabolism of glucose and glutamine to the TCA cycle (fig. S2C and S3A-D). Instead, 6-TG enhanced the contribution of glutamine-dependent reductive carboxylation, a well-known metabolic feature of cells with ETC defects(*21–23*) (fig. S3C,D). None of these proteomic and metabolic alterations were observed in cells lacking NUDT5 (fig. S2C and S3A-D), indicating that 6-TG’s effects on mitochondrial function also require NUDT5.

### NUDT5 regulates cellular thiopurine sensitivity differently from HPRT1

To more rigorously examine whether NUDT5 influences HPRT1-mediated purine salvage, we assessed this pathway by conducting stable isotope tracing using ^15^N-labeled hypoxanthine ([^15^N_4_]hypoxanthine). In the presence of this tracer, HPRT1 transfers four ^15^N nuclei into purine nucleotides, which can be detected by mass spectrometry as m+4 isotopologues (Fig. 2A). As expected, depletion of HPRT1 eliminated purine nucleotide labeling from [^15^N_4_]hypoxanthine (Fig. 2B and fig. S4A). However, WT and NUDT5-null cells displayed comparable levels of labeled purine nucleotides, indicating that NUDT5 is not required for purine salvage (Fig. 2B and fig. S4A). Furthermore, WT and NUDT5-null cells are less sensitive than HPRT1-null cells to inhibitors of DNPB, such as lometrexol (LTX) and methotrexate (MTX) (Fig. 2C and fig. S4B), consistent with persistent purine salvage. Direct measurement of HPRT1 enzymatic activity in lysates from NUDT5-null cells expressing empty vector, WT NUDT5, or NUDT5^E112Q^ revealed no differences (Fig. 2D). These data make it unlikely that 6-TG resistance of NUDT5-null cells results from a failure to convert 6-TG to 6-TGMP through salvage. To confirm this, we measured the relevant metabolites after 6-TG treatment. Intracellular 6-TG was barely detectable in WT and NUDT5-null cells after 24 hours of 6-TG treatment (Fig. 2E and fig. S4C). This was not due to defective uptake of 6-TG, but rather to its conversion to 6-T-GMP and subsequently 6-T-GTP and 6-thioguanosine, all of which were easily detected (Fig. 2F and fig. S4D). In contrast, HPRT1-null cells accumulated intracellular 6-TG but did not produce 6-T-GMP, 6-T-GTP, or 6-thioguanosine (Fig. 2E,F and fig. S4C,D). Taken together, these data indicate that NUDT5 supports thiopurine toxicity through a mechanism distinct from HPRT1-mediated purine salvage.

**Figure 2.**
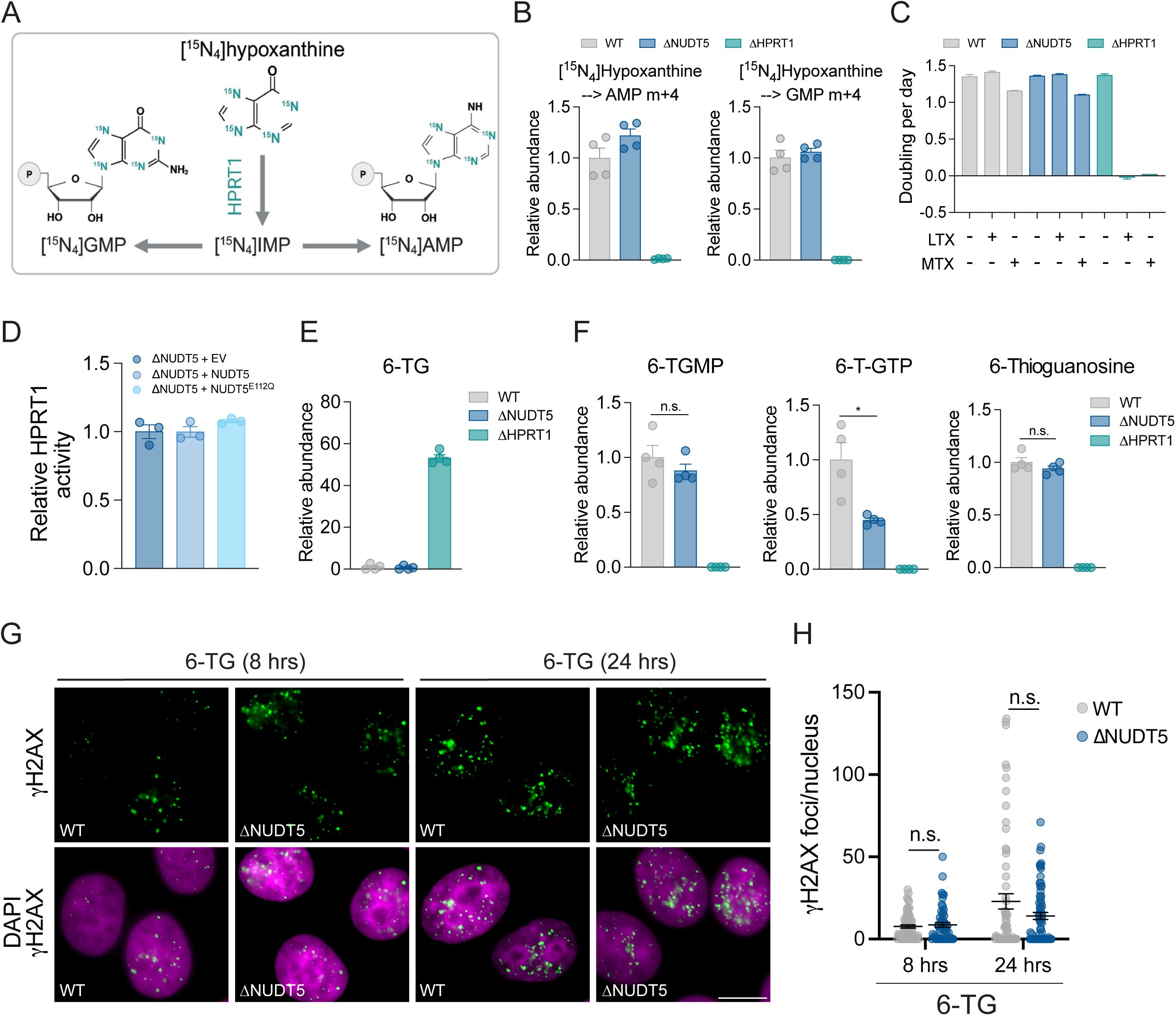
Loss of NUDT5 confers thiopurine resistance without suppressing purine salvage. **A.** Schematic illustrating labeling of purine nucleotides from [^15^N_4_]hypoxanthine. **B.** Relative abundance of m+4 AMP and m+4 GMP from [^15^N_4_]hypoxanthine in WT, ΔNUDT5, and ΔHPRT1 HeLa cells during 4 hours of tracing. Data are means from four replicates. **C.** Growth rates of WT, ΔNUDT5, and ΔHPRT1 HeLa cells treated with DMSO, 1 µM lometrexol (LTX), or 1 µM methotrexate (MTX). Data are from one of three independent experiments. **D.** Relative HPRT1 enzymatic activity in lysates from ΔNUDT5 HeLa cells that express empty vector (EV), WT NUDT5, or NUDT5^E112Q^. Data are means from three replicates. **E-F.** Relative abundance of 6-TG (**E**) and 6-T-GMP, 6-T-GTP, and 6-thioguanosine (**F**) in WT, ΔNUDT5, and ΔHPRT1 HeLa cells treated with 0.5 µg/mL 6-TG for 24 hours. Data are means from four replicates. **G.** Analysis of γH2AX foci in WT and ΔNUDT5 HeLa cells treated with 0.5 µg/mL 6-TG at the indicated time points. Nuclei were labeled with DAPI (magenta) and γH2AX foci are green. The scale bar represents 10 µm. **H.** Quantification of γH2AX foci per nucleus. Unpaired, two-sided t tests (**F**) and Two-Way ANOVA (**H**) were used for statistical analyses *: P < 0.05; n.s.: P > 0.05. Error bars denote SEM. BioRender was used to generate the illustration in **A**.

### NUDT5 is dispensable for 6-TG-induced DNA damage

Incorporation of thioguanines into the genome causes DNA mismatches and subsequent DNA breaks(*24–26*). To investigate whether NUDT5 loss mitigates DNA damage or enhances DNA repair to improve cell survival in the presence of thiopurines, we stained cells for γH2AX, a nuclear DNA damage marker. WT and NUDT5-null cells exhibit comparable numbers γH2AX foci after 6-TG treatment, indicating that NUDT5 loss does not confer resistance to thiopurines by blocking DNA damage (Fig 2G,H). Additionally, NUDT5 ablation did not protect cells against ionizing radiation (IR) or the DNA damaging agents cisplatin and etoposide (fig. S5A-C), suggesting that NUDT5 loss does not enhance DNA repair responses in general. Although previous studies showed that DNA mismatch repair (MMR) deficiency rendered colorectal cancer cell lines resistant to 6-TG(*24, 27*), MMR genes did not emerge as hits in the CRISPR screen data we analyzed (fig. S6A). To rule out the involvement of MMR in our model, we deleted the gene encoding MutL protein homolog 1 (MLH1), a key protein in the MMR pathway (fig. S6B). MLH1 depletion provided no protection against 6-TG treatment (fig. S6C). Therefore, NUDT5 specifically regulates responses to purine analogs, highlighting its potential role in regulating purine metabolism.

### Thiopurines suppress DNPB in a NUDT5-dependent fashion

Although DNA damage at least partially contributes to cell growth arrest and apoptosis after treatment with thiopurines, previous studies showed that thiopurine-induced cell death may also stem from suppression of DNPB, because accumulated thiopurine nucleotides inhibit phosphoribosyl pyrophosphate admidotransferase (PPAT), the rate-limiting enzyme in DNPB(*1, 15*). We used metabolomics and isotope tracing to characterize the effects of 6-TG on cellular metabolism. In WT cells, 6-TG induced marked changes in the metabolome (fig. S7A). Metabolite set enrichment analysis identified purine metabolism as one of the most significantly altered pathways, with many purine-associated metabolites reduced following 6-TG treatment (Fig. 3A,B). Additionally, isotope tracing with [U-^13^C]glucose revealed that 6-TG suppresses labeling of purine nucleotides, indicating reduced purine synthesis (Fig. 3C and S7B). However, depletion of NUDT5 essentially eliminated the effect of 6-TG on purine labeling (Fig. 3C). Suppressed purine nucleotide synthesis in WT cells was not due to decreased synthesis of R5P, which was highly labeled under all conditions (fig. S7C). Instead, 6-TG induced an accumulation of labeled R5P and other metabolites related to the pentose phosphate pathway (PPP) (fig. S7D-F).

**Figure 3.**
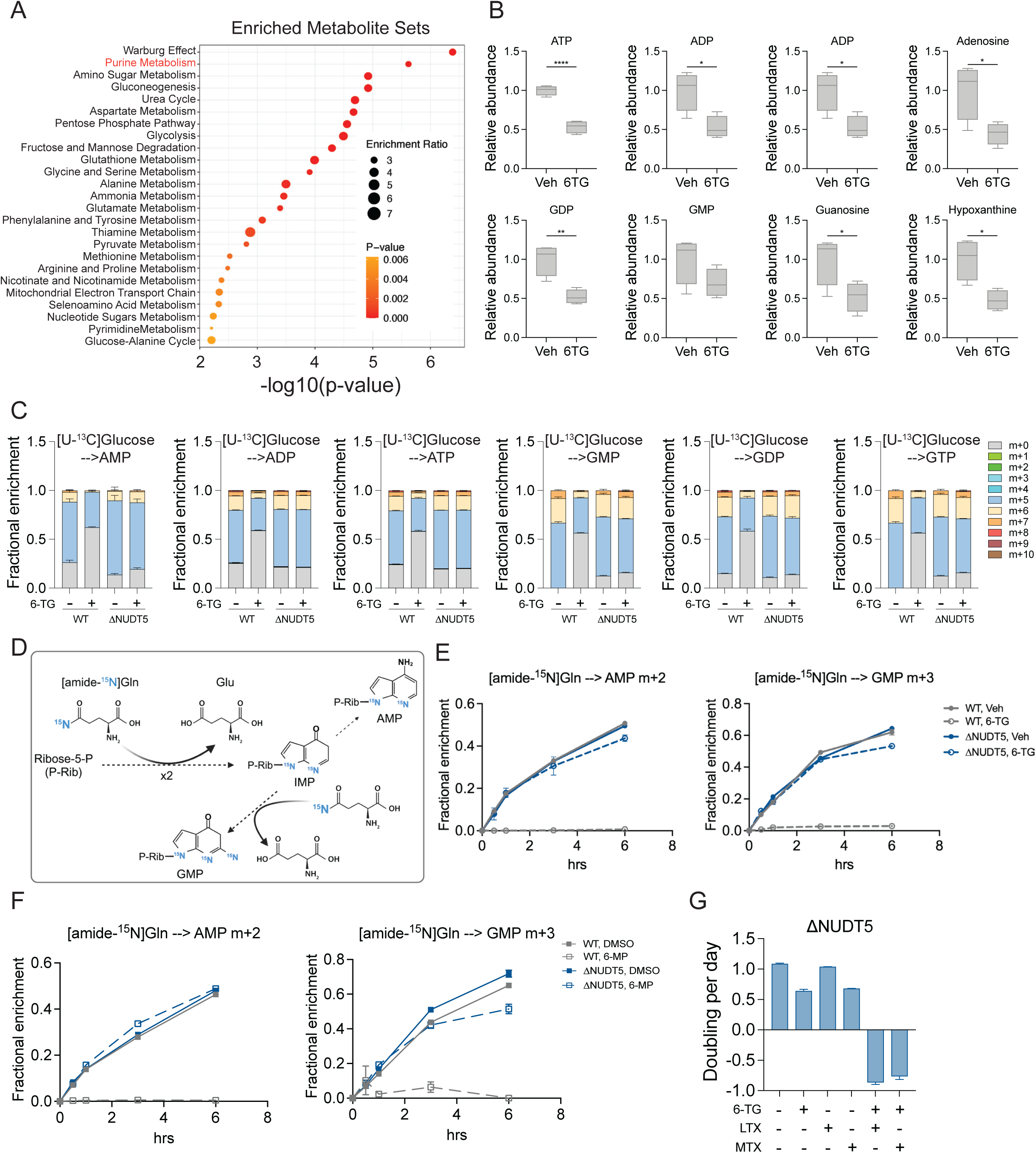
NUDT5 ablation overcomes the thiopurine-induced block in *de novo* purine nucleotide synthesis. **A.** Metabolite set enrichment analysis comparing vehicle and 24 hours of 0.5 µg/mL 6-TG-treated HeLa cells. **B.** Relative abundance of the indicated purine metabolites in vehicle or 0.5 µg/mL 6-TG-treated HeLa cells. Data are means from four replicates. **C.** ^13^C labeling in the indicated purine nucleotides after 6 hours of culture with [U-^13^C]glucose in WT and ΔNUDT5 HeLa cells pre-treated with vehicle or 0.5 µg/mL 6-TG for 24 hours. **D.** Schematic illustrating labeling of purine nucleotides from [amide-^15^N]glutamine. **E-F.** Time-dependent fractional enrichment of m+2 AMP and m+3 GMP from [amide-^15^N]glutamine in WT and ΔNUDT5 HeLa cells pre-treated with 0.5 µg/mL 6-TG (**E**) or 20 µM 6-MP (**F**) for 24 hours. Data are means from three replicates. **G.** Growth rates of ΔNUDT5 HeLa cells treated with 0.5 µg/mL 6-TG, 1 µM LTX, 1 µM MTX or combinations. Unpaired, two-sided t tests were used for the statistical analyses (**B**) ****: P < 0.0001; **: P < 0.01; *: P < 0.05. Error bars denote SEM. BioRender was used to generate the illustration in **D**.

To more precisely assess DNPB activity, we cultured cells with [amide-^15^N]glutamine, which delivers ^15^N to purine nucleotides largely through DNPB (Fig. 3D). Strikingly, thiopurines nearly abolished DNPB in WT cells but had no effect in NUDT5-null cells (Fig. 3E-3F), indicating that thiopurines suppress DNPB in a NUDT5-dependent fashion. However, basal DNPB in the absence of 6-TG was not affected by either knockout or overexpression of NUDT5 (Fig. 3E-3F and fig. S7G). Of note, while thiopurines such as 6-MP have been documented to impair purine salvage, presumably as competitive inhibitors(*28*), the doses used here instead modestly enhanced the contribution of purine salvage to AMP and GMP in WT cells (fig. S8A,B), likely as a compensatory response to the drastic reduction in DNPB.

Native purine nucleotides are essential for DNA repair, replication, transcription, and other biological activities to support cell growth and proliferation. Genotoxic insults induced by thiopurines may increase the demand for purine nucleotides, which basal rates of purine salvage alone cannot meet under conventional cell culture conditions. Congruently, supplementation with excess purine nucleosides or bases to enhance purine salvage rescued WT cell growth in the presence of 6-TG (fig. S8C). Furthermore, while NUDT5-null cells can partially tolerate either thiopurines or DNPB inhibitors individually, they are highly sensitive to concomitant treatment (Fig. 3G and fig. S8D). Our results indicate that loss of NUDT5 enables persistent DNPB activity during thiopurine treatment.

### NUDT5 facilitates the disassembly of purinosomes in response to purine salvage stimulation

Since enhanced purine salvage represses DNPB by increasing cellular AMP and GMP, we hypothesized that NUDT5 also engages in DNPB suppression in response to elevated native purine nucleotides. To test this, we pre-treated the cells with hypoxanthine and inosine to stimulate purine salvage prior to assessing DNPB using [amide-^15^N]glutamine tracing. As expected, pretreatment of either hypoxanthine or inosine caused a dramatic reduction in DNPB, which was alleviated by NUDT5 loss (Fig. 4A).

**Figure 4.**
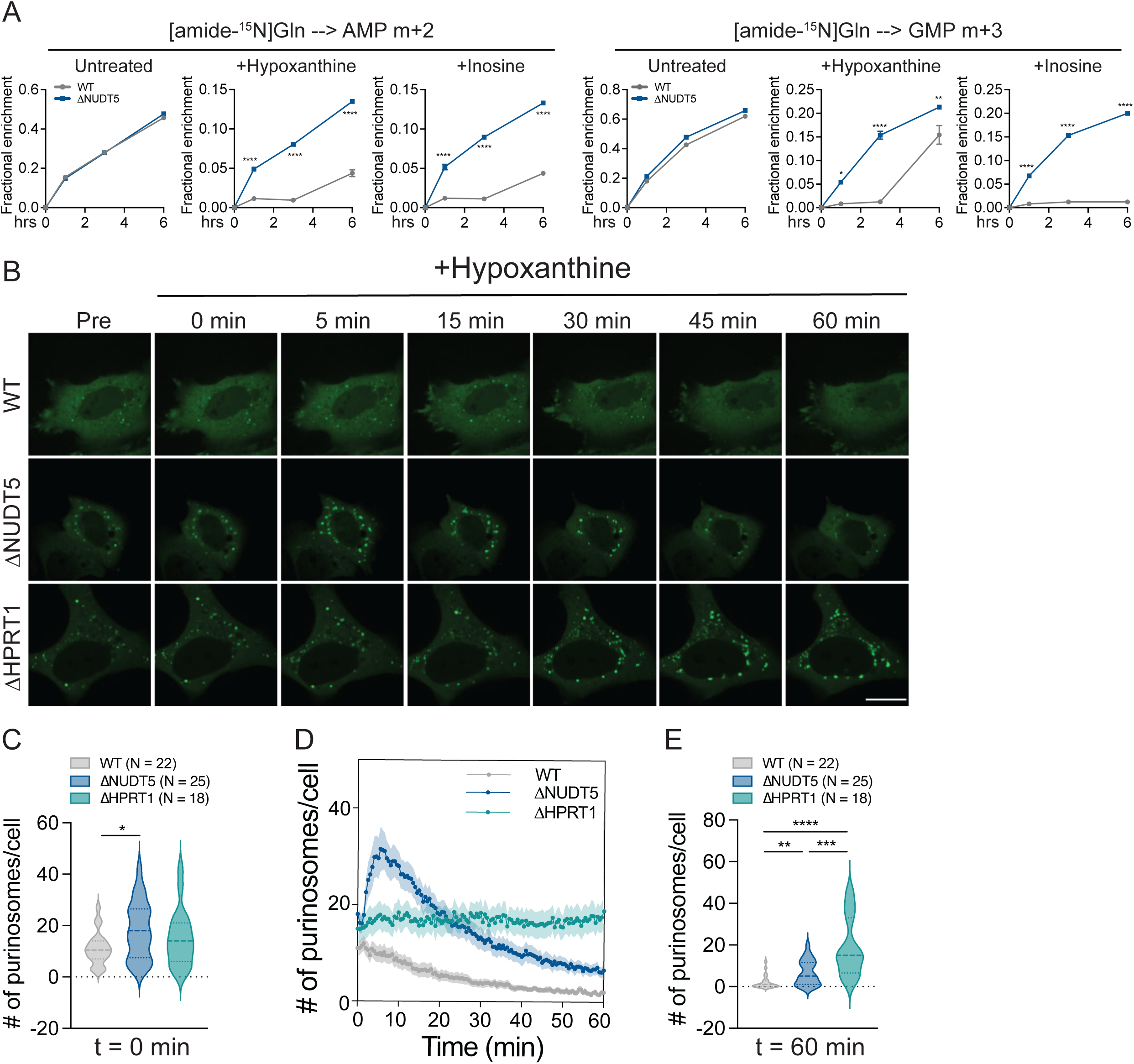
NUDT5 facilitates purinosome disassembly during purine salvage. **A.** Time-dependent fractional enrichment of m+2 AMP and m+3 GMP from [amide-^15^N]glutamine in WT and ΔNUDT5 HeLa cells pre-treated with or without 20 µM hypoxanthine or inosine for 24 hours. Data are means of three replicates. **B.** Time-lapse imaging showing purinosome dispersal upon treatment with 20 µM hypoxanthine at t=0 min in WT, ΔNUDT5 and ΔHPRT1 HeLa cells. The fluorescence signals are from GFP-tagged FGAMS. The scale bar represents 10 µm. **C.** Quantification of purinosome count in (**B**) at t=0. **D.** Quantification of purinosome count over one hour after hypoxanthine supplementation in (**B**). **E.** Quantification of purinosome count in (**B**) at t=60 minutes. Two-way ANOVA (**A**) and One-way ANOVA (**C** and **E**) were used for statistical analyses. ****: P < 0.0001; ***: P < 0.001; **: P < 0.01; *: P < 0.05.

Next, we explored the mechanism by which NUDT5 suppresses DNPB when purine salvage is active. Although PPAT is subject to allosteric inhibition, in vitro analyses using purified PPAT showed that supraphysiological levels of AMP or GMP are required to achieve such inhibition(*29–33*). Therefore, in intact cells, other factors may be involved in regulating allosteric inhibition of PPAT or other aspects of DNPB. Over a decade ago, the Benkovic group reported that PPAT and several other DNPB enzymes form a subcellular structure called the purinosome that promotes metabolite channeling and accommodates labile intermediates such as PPAT’s product, 5-phosphoribosyl-1-amine (PRA), which has a half-life of 38 seconds at 37°C and pH 7.5(*9–11, 13, 34*). Purinosomes are dynamically regulated and reversible. They appear under conditions of purine deprivation, then quickly disperse upon supplementation with salvageable purine bases(*9*). Therefore, dissociation of purinosomes in response to robust purine salvage is linked to the negative regulation of DNPB. To test whether NUDT5 modulates purinosome assembly and disassembly, we expressed GFP-tagged phosphoribosylformyl glycinamidine synthase (FGAMS), a core purinosome protein, in WT, NUDT5-null and HPRT1-null cells. These cell lines were cultured in purine-depleted medium to induce purinosome formation, indicated by clustering of FGAMS-GFP into punctate structures(*9, 12, 13*) (Fig. 4B). Ablation of NUDT5 modestly increased baseline purinosome formation (Fig. 4C), consistent with the [amide-^15^N]glutamine tracing results showing comparable DNPB between WT and NUDT5-null cells (Fig. 3E,F). We next performed live cell imaging to monitor purinosome kinetics and quantify the number of FGAMS-GFP puncta in cells during one hour after hypoxanthine repletion (Fig. 4D). As previously reported, upon supplementation with hypoxanthine, punctate FGAMS-GFP structures in WT cells completely disappeared within about one hour(*9*)(Figures 4B-E, Movie 1). Also as expected, hypoxanthine did not disperse punctate FGAMS-GFP in HPRT1-null cells (Fig. 4B-E, Movie 2). However, in NUDT5-null cells, addition of hypoxanthine initially stimulated punctate FGAMS-GFP signal, followed by a delay in dispersal, resulting in substantial residual punctate signal after one hour (Fig. 4B-E, Movie 3). Taken together, these data indicate that to respond to purine abundance and suppress DNPB, NUDT5 promotes purinosome disassembly.

### NUDT5 interacts with PPAT to regulate DNPB in response to purine salvage stimulation

To elucidate how NUDT5 regulates purinosome dynamics, we analyzed the NUDT5 interactome using BioPlex 3.0(*35, 36*). Surprisingly, we found that NUDT5 interacts with PPAT, a core purinosome component (Fig. 5A). Previous work showed that NUDT5 primarily functions as a dimer whereas PPAT forms either dimers or tetramers(*18, 37, 38*). Our in silico screen of human protein-protein interactions, which combines coevolution analysis and deep learning methods, also identified the interaction between NUDT5 and PPAT(*39*). Thus, we modeled the interaction of NUDT5 as a dimer and PPAT as a tetramer by AlphaFold Server (AF3)(*40*). This analysis revealed that two NUDT5 dimers can form a complex with a PPAT tetramer (fig. S9A). We then mapped the PPAT-NUDT5 interaction sites and identified Arginine 70 (R70) in NUDT5 as one of the key residues for this interaction (Fig. 5B). To test whether the PPAT-NUDT5 interaction is important for DNPB suppression by purine supplementation, we ectopically expressed NUDT5 harboring an arginine 70 to alanine mutation (NUDT5^R70A^) in NUDT5-null cells. Like WT NUDT5, but not catalytically inactive NUDT5 (NUDT5^E112Q^), NUDT5^R70A^ diminished cellular ADPR levels, indicating that its enzymatic function is intact (fig. S9B). However, unlike WT NUDT5 and NUDT5^E112Q^, NUDT5^R70A^ failed to associate with PPAT in an immunoprecipitation assay (Fig. 5C). Remarkably, disrupting the PPAT-NUDT5 interaction in this manner hinders purinosome disassembly (Figures 5D,E, S9C, Movies 4-7) and mitigates DNPB suppression by excessive supplementation with hypoxanthine or inosine (Fig. 5F,G).

**Figure 5.**
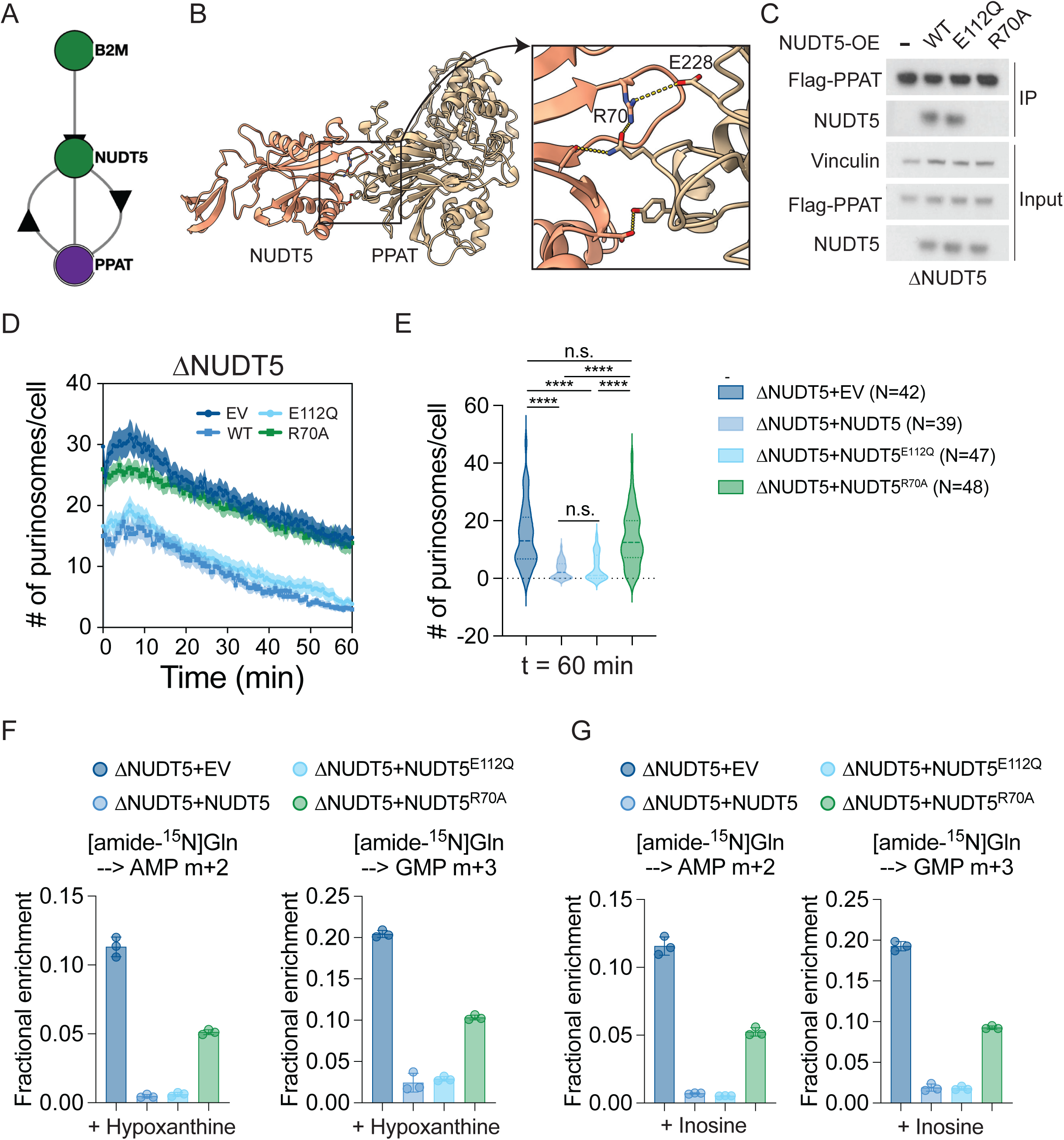
NUDT5 interacts with PPAT to regulate *de novo* purine synthesis. **A.** Bio-plex data showing the interaction between PPAT and NUDT5. **B.** PPAT-NUDT5 interaction depicted by AlphaFold3. **C.** Western blot showing the interaction between Flag-PPAT and WT NUDT5 or NUDT5^E112Q^ but not NUDT5^R70A^. Vinculin is the loading control for the input samples. **D.** Quantification of purinosome count in ΔNUDT5 HeLa cells that express empty vector (EV), WT NUDT5, NUDT5^E112Q^, or NUDT5^R70A^ during the hour after 20 µM hypoxanthine treatment. **E.** Quantification of purinosome count (**D**) at t=60 minutes. **F-G.** Fractional enrichment of m+2 AMP and m+3 GMP from [amide-^15^N]glutamine in ΔNUDT5 HeLa cells that express empty vector (EV), WT NUDT5, NUDT5^E112Q^, or NUDT5^R70A^ during 4 hours of tracing. The cells were treated with 20 µM hypoxanthine (**F**) or inosine (**G**) for 24 hours. Data are means from three replicates. One-way ANOVA was used for statistical analysis (**E**). ****: P < 0.0001; n.s.: P > 0.05. Error bars denote SEM.

We then used the standalone version of AlphaFold3, available on GitHub (https://github.com/google-deepmind/alphafold3), to investigate the interaction between PPAT and its small-molecule ligands, including AMP, GMP, and PRPP. Unlike the online AlphaFold server, which limits users to a predefined set of small molecules (such as AMP), the standalone version allows the modeling of a broader range of small molecules, including GMP and PRPP. Superimposition of PPAT’s structure in its AMP-, GMP- or PRPP-bound state to its structure in complex with NUDT5 showed no major conformational changes, with root-mean-square-deviation (rmsd) values of 2.158, 2.049, and 1.477 angstroms for the AMP, GMP, and PRPP-bound forms, respectively (fig. S10A-C). This analysis implies no substantial effect of NUDT5 on PPAT’s binding capacity to AMP, GMP or PRPP.

### Disrupting PPAT-NUDT5 interaction is sufficient to confer resistance to thiopurines

Cells expressing WT or NUDT5^E112Q^ suppress DNPB upon exposure to thiopurines, whereas cells expressing NUDT5^R70A^ display comparable DNPB under these conditions (Fig. 6A and fig. S11A). Since perturbation of the PPAT-NUDT5 interaction counteracts thiopurine-induced DNPB suppression, we reasoned that the loss of the interaction would also allow cells to survive thiopurine treatment. Indeed, compared to cells expressing either WT or NUDT5^E112Q^, cells expressing NUDT5^R70A^ maintained growth and resisted apoptosis when treated with thiopurines, similar to NUDT5-null cells (Fig. 6 B-D and fig. S11B). These data substantiate the conclusion that the NUDT5’s ability to interact with PPAT, not its enzymatic activity, regulates DNPB and thiopurine resistance through modulation of purinosome disassembly upon salvage of (thio)purines (fig. S11C).

**Figure 6.**
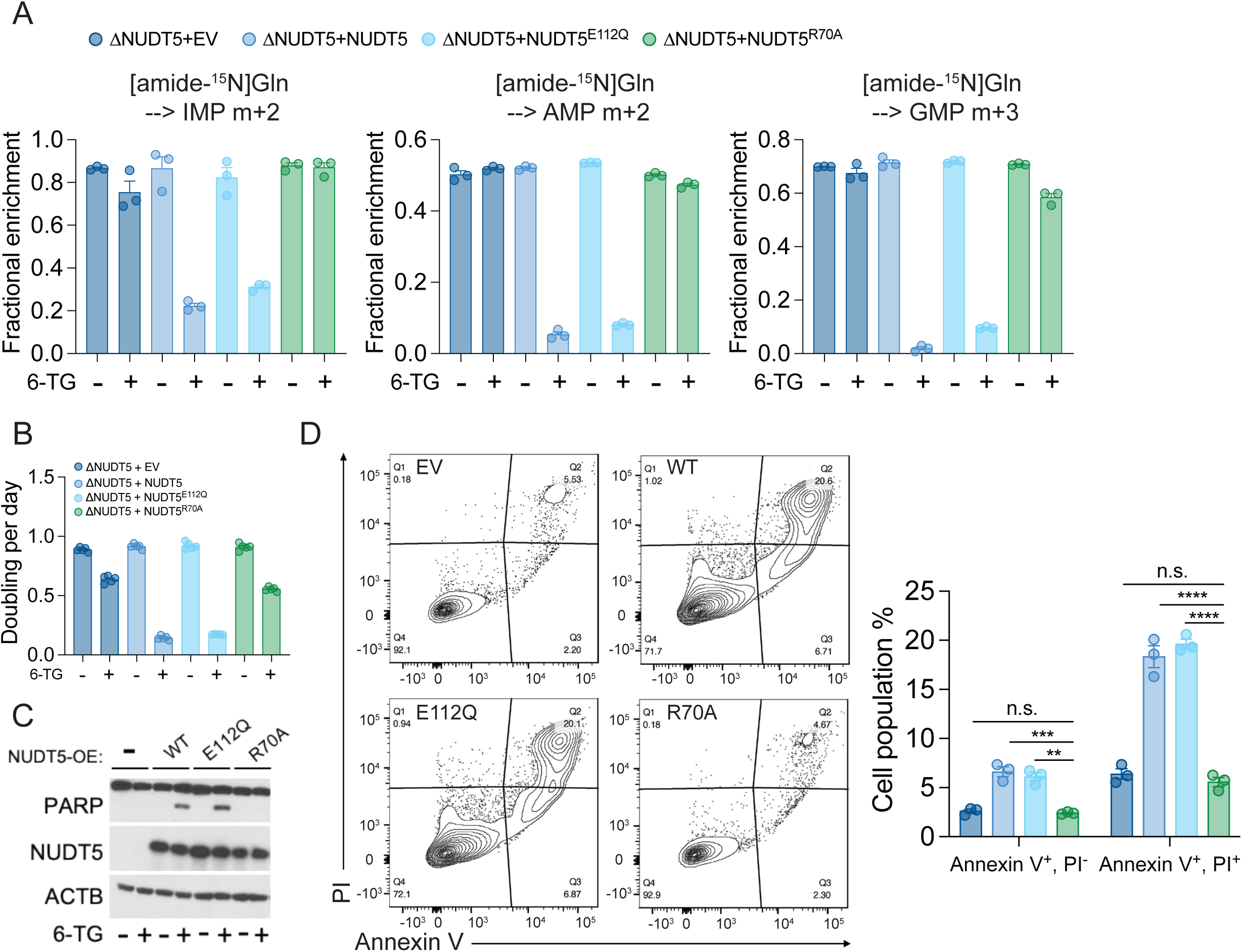
Disruption of the NUDT5-PPAT interaction induces thiopurine resistance. **A.** Fractional enrichment of m+2 IMP, m+2 AMP and m+3 GMP from [amide-^15^N]glutamine in ΔNUDT5 HeLa cells that express empty vector (EV), WT NUDT5, NUDT5^E112Q^, or NUDT5^R70A^ during 6 hours of tracing. The cells were pre-treated with 0.5 µg/mL 6-TG for 24 hours. Data are means from three replicates. **B.** Growth rates of ΔNUDT5 HeLa cells that express empty vector (EV), WT NUDT5, NUDT5^E112Q^, or NUDT5^R70A^, treated with 2 µg/mL 6-TG. Data are from one of three experiments. **C.** Western blot assessing cleaved PARP in ΔNUDT5 HeLa cells that express empty vector (EV), WT NUDT5, NUDT5^E112Q^, or NUDT5^R70A^, treated with 0.5 µg/mL 6-TG for 24 hours. β-actin (ACTB) is the loading control. **D.** Apoptosis analysis in ΔNUDT5 HeLa cells that express empty vector (EV), WT NUDT5, NUDT5^E112Q^, or NUDT5^R70A^, treated with 0.5 µg/mL 6-TG for 48 hours. To the left are representative contour plots. To the right are bar graphs showing percentage of apoptotic (Annexin V+, PI −) and dead (Annexin V+, PI+) cells. Data are means from three replicates. Two-way ANOVA was used were used for statistical analysis (**D**). ****: P < 0.0001; ***: P < 0.001; **: P < 0.01; n.s.: P > 0.05. Error bars denote SEM.

## Discussion

Thiopurines are used to treat ALL and inflammatory diseases such as inflammatory bowel disease and rheumatoid arthritis(*41, 42*). These drugs have substantial systemic toxicity, including bone marrow suppression, hepatotoxicity, pancreatitis, and gastrointestinal intolerance(*43*). Given that thiopurines require metabolic processing to exert both their therapeutic and toxic effects, human genetic variants that affect thiopurine metabolism impact clinical responses to treatment(*44–47*). For example, patients with *HPRT1* variants that impair purine salvage are refractory to thiopurine therapy(*46*), whereas loss-of-function mutations in Nudix Hydrolase 15 (*NUDT15*), a key enzyme responsible for detoxifying thioguanine nucleotides, leads to increased sensitivity to thiopurines(*45, 48*). In this study, we identified *NUDT5* as a new component in cellular responses to thiopurines. Our work is consistent with recent reports describing NUDT5’s involvement in DNPB(*49–51*).

Unlike HPRT1, NUDT15 and thiopurine methyltransferase (TPMT), we found that NUDT5 regulates thiopurine responses independently of its known enzymatic function(*44–46*). Instead, NUDT5 binds PPAT to suppress DNPB when exposed to thiopurines. Disrupting the PPAT-NUDT5 interaction through mutation of a NUDT5 residue at the interface with PPAT enables persistent DNPB activity during 6-TG treatment. This allows cells to resist the effects of the drug, possibly by continuously providing purine nucleotides for DNA repair and to preserve cellular homeostasis. Our findings highlight a novel regulatory mechanism of NUDT5 on thiopurine response. Clinical evaluation of NUDT5 expression may be beneficial to personalize thiopurine doses.

Beyond its role in thiopurine-induced cell death, the PPAT-NUDT5 interaction is critical for preventing simultaneous activation of DNPB and purine salvage, contributing to optimal cellular purine nucleotide levels. It is well established that inhibition of DNPB activates purine salvage and *vice versa*, reflecting a metabolic plasticity by which cells ensure a continuous supply of purine nucleotides(*3, 8*). However, properly limiting either one of these pathways is important, because excessive purine nucleotide production can disrupt pyrimidine-purine balance, causing DNA replication stress and cell cycle arrest(*5*). Excess purine accumulation through chronic hyperactivation of DNPB is also deleterious, causing hyperuricemia and associated toxicities in patients. In line with this, active purine salvage suppresses DNPB, and our findings further elucidate the molecular mechanism underlying this suppression.

So far, human PPAT has not been successfully purified. Studies on purified PPAT from non-mammalian organisms revealed that AMP and GMP allosterically inhibit PPAT in a dose-dependent manner(*29–31*). However, this inhibition requires millimolar concentrations that exceed physiological levels of AMP and GMP in both bacteria and human cells(*29–33*). Therefore, additional mechanisms must contribute to purine nucleotide-mediated allosteric inhibition of PPAT in intact cells and tissues. Protein-protein interactions and post-translational modifications are known to induce conformational changes in enzymes, thereby modulating their catalytic activities. Interestingly, previous work showed that the binding of AMP or GMP promotes human PPAT tetramerization whereas the PPAT activator PRPP keeps it in a dimeric state(*37*). Whether NUDT5 binding to PPAT influences its catalytic activity through alteration of PPAT oligomerization status is an appealing question for future studies.

In yeast, PPAT forms condensates through phase separation to enhance DNPB, a process that can be disrupted by elevated AMP and GMP levels(*33*). In mammalian cells, PPAT together with other DNPB enzymes assemble into purinosomes to facilitate efficient DNPB(*9*). Similar to yeast PPAT condensation, purinosome assembly and dispersal are dynamically regulated by purine nucleotide levels(*9*). Our work shows that NUDT5 modulates purinosome disassembly during purine salvage. This negative regulation is critical to prevent inappropriate DNPB and avoid the consequences of purine nucleotide overproduction. Taken together, our findings identify NUDT5 as a novel DNPB regulator to preclude excessive purine nucleotide synthesis, potentially governing purine-pyrimidine balance to maintain cellular homeostasis.

## Acknowledgements

We thank members of the DeBerardinis laboratory for helpful discussions. This article is subject to HHMI’s Open Access to Publications policy. HHMI lab heads have previously granted a nonexclusive CC BY 4.0 license to the public and a sublicensable license to HHMI in their research articles. Pursuant to those licenses, the author-accepted manuscript of this article can be made freely available under a CC BY4.0 license immediately upon publication. RJD is supported by the Howard Hughes Medical Institute Investigator Program, grants from the National Institutes of Health (R35CA220449, P50CA196516, P50CA070907), the Moody Foundation (Robert L. Moody, Sr. Faculty Scholar Award) and the Eugene McDermott Endowment for the Study of Human Growth and Development. PTN is supported by the National Institute of General Medical Sciences of the National Institutes of Health under Award Number K99GM151439. The CRI Metabolomics Facility is supported by funding from the Cancer Prevention Research Institute of Texas (CPRIT Core Facilities Support Award RP240494).

## Author Contributions

R.J.D., and Z.W., conceived and designed the study. Z.W. designed and conducted experiments; and analyzed the data. T.P.M., F.C. designed, optimized and validated the metabolomics methods. R.J.D. and Z.W. interpreted the data and wrote the manuscript. P.T.N., V.S., R.Y., T.D., J.C., Z.S., L.C., J.Z., M.D.M., I.A., X.L., L.G.Z., R.W., J.S.P., X.J., W.C., Q.C., and Y.Z., contributed to conducting experiments, data analysis, or interpretation.

## Methods

### Cell culture

HeLa cells were a gift from Dr. Javier Garcia-Bermudez’s laboratory at UT Southwestern. A549 cells were purchased from American Type Culture Collection (ATCC). Both cell lines were cultured in RPMI-1640 (Thermo Fisher Scientific, CB-40234) supplemented with 10% fetal bovine serum (FBS), at 37°C in a humidified atmosphere with 5% CO_2._ Cells were routinely subject to mycoplasma testing.

### Gene deletion and over-expression

To generate *HPRT1* and *NUDT5* knockouts with a Cas9 D10A nickase system, cells were transfected with two modified pDG461 constructs(*52*) with mGreenLantern and mScarlett fluorescent markers. Each vector contains two sgRNAs against the indicated genes or scrambled controls. The cells were sorted for GFP and RFP double-positive populations 48 hours after transfection. To generate cell lines that over-express WT or mutant NUDT5, the cells were transfected with constructs that contain cDNAs of the genes of interest integrated at the Rogi2 safe harbor locus(*53*). The cells were then selected by culturing with 1 µg/mL puromycin (Thermo Fisher Scientific, NC9138068) 48 hours after transfection until all non-transfected cells were dead. To generate sgScr and sgMLH1 HeLa cells, the indicated gRNAs were cloned into the LentiCRISPRv2 vector(*54*), a gift from Feng Zhang (Addgene plasmid # 52961; http://n2t.net/addgene:52961; RRID:Addgene_52961) followed by transfection into 293T cells using Lipofectamine 3000 (Thermo Fisher Scientific L3000015) at a ratio of 2:1 for psPAX2:pMD2G to generate lentivirus. To generate HeLa cells expressing streptavidin-binding Flag-Strep-PPAT, the cDNA was cloned into the PMXS-IRES-Bsd retroviral expression vector followed by transfection into 293T cells using Lipofectamine 3000 (Thermo Fisher Scientific L3000015) at a ratio of 2:1 for of Gag-Pol:VSVG to generate retrovirus. All virus-containing medium was collected 48 hours after transfection and filtered through 0.45µm membranes, then immediately added to HeLa cell cultures containing 4 µg/mL polybrene (Sigma, TR-1003-G). After 24 hours, 1 µg/mL puromycin and 10 µg/mL blasticidin (Thermo Fisher Scientific, NC1366670) were used to select sgScr/sgMLH1 expressing cells and Flag-Strep-PPAT overexpressing cells, respectively, until all non-infected cells were dead. Both deletion and overexpression of the proteins were validated by western blot. DNA oligos were purchased from IDT. The gRNA sequences were:

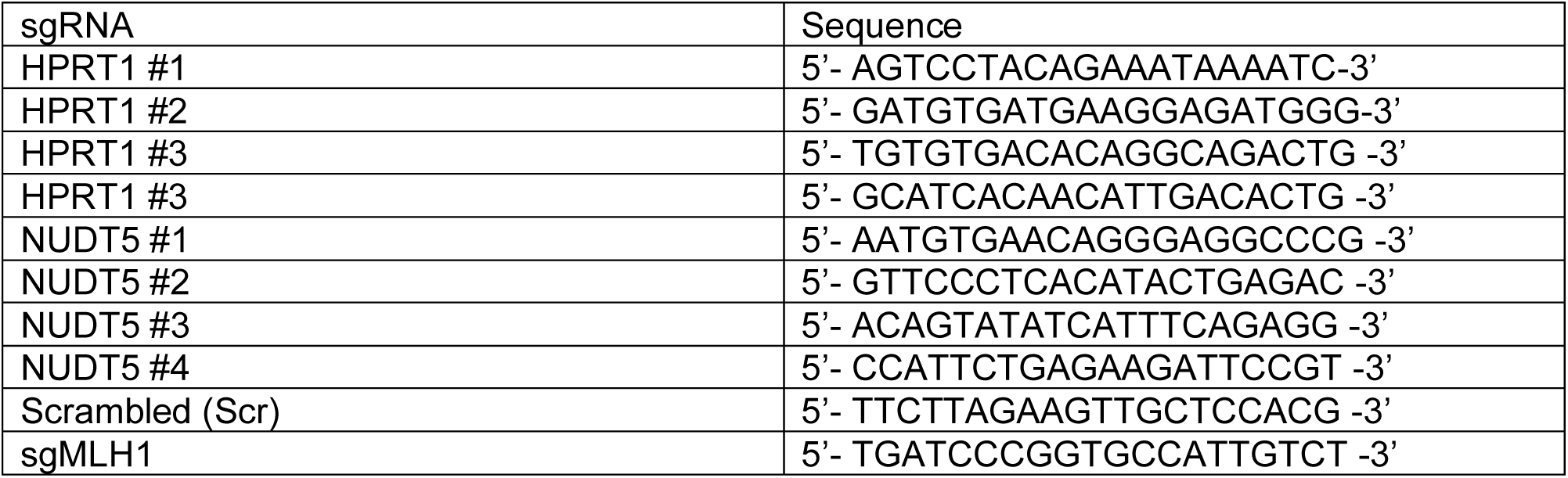

### Quantitative proteomics

WT and NUDT5 null cells were treated with vehicle or 0.5 µg/mL 6-TG for 24 hours. The cells were rinsed twice with ice-cold PBS then lysed with a solution containing 5% SDS in 50 mM TEAB with protease and phosphatase inhibitors. Cells were scraped on ice and transferred to 1.5 mL Eppendorf tubes and incubated at room temperature for 15 minutes to allow complete lysis. Proteins were quantified using DC Protein Assay Kit (Bio-Rad, 5000111) and normalized to the same concentration. These samples were subjected to overnight digestion with trypsin using S-Trap (Protifi) after disulfide bond reduction and alkylation. The eluted peptide from the S-Trap was dried and reconstituted in 100 mM TEAB buffer. The samples were labeled using a TMT10plex Isobaric Mass Tagging Kit (Thermo) following the manufacturer’s instructions. The combined sample was passed through solid-phase extraction using an Oasis HLB plate (Waters) and dried in a SpeedVac. The dried sample was reconstituted in 0.1% TFA buffer containing 2% acetonitrile, and then diluted down to ∼1 ug of peptides before injection. A Thermo Orbitrap Eclipse MS system coupled to an Ultimate 3000 RSLC-Nano liquid chromatography system was used to analyze the peptides in the UT Southwestern Proteomics Core Facility, as previously described(*8*).

To remove potential bias from differing sample intensities, normalization factors were calculated by summing abundances across non-missing entries and dividing by the mean total intensity. These normalization factors were used to scale each dataset accordingly. Statistical analyses were performed using a custom function based on linear modeling and t-tests. For each analyte, we assessed: 1) treatment effect within genotypes using Welch’s t-test and Cohen’s d effect size; 2) genotype effect under each treatment using similar t-tests; and 3) interaction effects using two-way ANOVA models of the form value ∼ genotype * treatment. P-values were adjusted using the Benjamini–Hochberg method for false discovery rate (FDR) control. Enrichment of pathway terms was performed using a hypergeometric test to assess the overlap between curated pathway gene sets from public databases Wikipathways of significantly altered proteins, selected by fold-change direction and adjusted p-value < 0.05). Z-scored expression values were used for heatmap visualization to compare selected features across genotypes, treatments, and replicates. Data were visualized using the ComplexHeatmap package(*55*).

### Targeted metabolomics

To extract metabolites, the cells were rinsed in ice-cold saline and quenched with 80% acetonitrile. The cells were incubated at −80°C for at least 20 minutes, then transferred onto ice, scraped and subjected to three freeze-thaw cycles between liquid nitrogen and a 37°C water bath. Afterwards, the samples were vortexed for 1 minute and spun down at 4°C at 20,160 x g for 15 minutes. The supernatants were collected for the second spin-down under the same condition. Then, the supernatants were transferred into fresh Eppendorf tubes followed by protein quantification and normalization prior to analysis on Thermo Exploris or Q-Exactive liquid chromatograph/mass spectrometry systems. Chromatographic separation of metabolites was performed using a Vanquish UHPLC system equipped with a ZIC-pHILIC column (Millipore-Sigma, Burlington, MA) as previously described(*8*). Extracted ion chromatograms (XICs) were produced with a mass tolerance of 5 ppm. Analyte identities were verified using purified standards and product ion spectra. MetaboAnalyst 6.0 was used to analyze the metabolomics data and generate the principal component analysis plot(*56*).

### Gas chromatography/mass spectrometry (GC/MS)

To extract metabolites, the cells were rinsed with cold saline, scraped and subjected to three freeze-thaw cycles as above. After vortexing for 1 minute, the samples were centrifuged at 20,160 x g at 4°C for 15 min. The supernatants were transferred to new Eppendorf tubes and dried in a SpeedVac concentrator overnight. The dried metabolites were re-suspended in 30 µL of anhydrous pyridine containing 10 mg/mL methoxyamine. After a short vortex and centrifugation, the supernatants were transferred to GC/MS autoinjector vials and incubated at 75°C for 15 minutes. 70 µL N-(tert-butyldimethylsilyl)-N-methyltrifluoroacetamide (MTBSTFA) was added into each vial followed by incubation at 75°C for 1 hour. 1 µL of each sample was injected onto an Agilent 5973N or 5975C Mass Spectrometer coupled to Agilent 6890 or 7890 gas chromatographs. EL-MAVEN was used to analyze the data and MATLAB was used to correct for natural abundance as previously described(*57*).

### Stable isotope tracing

For [U-^13^C]glucose tracing, cells were cultured in glucose-free RPMI medium (Sigma, R1383-L) supplemented with 11 mM [U-^13^C]glucose (Cambridge Isotope Laboratories, CLM481-0.25) and 10% dialyzed FBS (Gemini Bio-Products, 100108). For glutamine tracing, cells were cultured in glutamine-free RPMI medium (Sigma, R0883) supplemented with 2 mM [amide-^15^N]glutamine (Cambridge Isotope Laboratories, NLM-557-1) or 2 mM [U-^13^C]glutamine (Cambridge Isotope Laboratories, CLM-1822-0) and 10% dialyzed FBS. For [^15^N_4_]hypoxanthine tracing, cells were cultured in RPMI medium containing 10% dialyzed FBS and 10 µM [^15^N_4_]hypoxanthine (Cambridge Isotope Laboratories, NLM-8500-0.1). If cells were pretreated with any drugs, they were exposed to the same drugs during tracing. Tracing time points were indicated in the figure legend. Extracted metabolites were subject to analyses with GC/MS or Q-Exactive MS as described above, or with an AB SCIEX QTRAP 5500 LC/triple quadrupole MS (Applied Biosystems SCIEX) as previously described(*8*).

### HPRT1 enzymatic activity analysis

HPRT1 enzyme activity was measured using the Precise HPRT1 assay kit (Novo CIB, K0709-01-2) according to the manufacturer’s instructions. Briefly, HeLa cells were cultured in 15 cm plates, washed once with PBS, collected by scraping, and lysed in ice-cold lysis buffer containing 150 mM NaCl, 10 mM Tris-HCl pH 7.4, 1 mM EDTA, and 1% NP-40. The lysates were centrifuged at 18,000 x g for 10 minutes at 4°C. Protein concentrations were determined using the DC Protein Assay Kit, then the samples were diluted to the same concentration. Each enzymatic reaction included 5 µL of sample or positive control (human recombinant HPRT enzyme) and 100 µL of the reaction mixture containing NAD, DTT, and bacterial IMPDH. The reactions were performed either with or without 2 mM PRPP at 37°C. Absorbance at 340 nm was measured every 2 minutes for 2 hours.

### Cell doubling analysis

Cells were seeded in clear, flat 96 well plates the day before treatment. To count the cell number, we stained the cells with 1 µg/mL Propidium Iodide (PI) (Thermo Fisher Scientific, P3566) and 5 µg/mL Hoechst (Thermo Fisher Scientific, 62249) in PBS at 37°C for at least 15 minutes. The plates were then analyzed using a Celigo Imaging Cytometer. Live cell numbers were calculated by subtracting the PI-positive cells from the Hoechst-positive cells. The doubling rates were calculated as previously described(*58*).

### Immunoblotting

Immunoblotting was performed as previously described (*8*). In brief, RIPA buffer (Boston BioProducts, BP-115) containing proteinase and phosphatase inhibitors (Thermo Fisher Scientific, 78444) was used to lyse cells. Samples were spun down at 20,160 x g at 4°C for 10 minutes followed by protein quantification using the DC Protein Assay Kit. Equal amounts of protein were loaded on the gel, followed by transfer to PVDF membranes (Thermo Fisher Scientific, 88518). Membranes were soaked in methanol for a few seconds and then rinsed with DI water prior to air drying. The dried membrane was incubated with primary antibodies diluted in filtered 5% BSA in PBS with 0.1% Tween-20 (PBST) overnight in the cold room. The membranes were washed with PBS 3 times for 5 minutes at room temperature followed by incubation with the secondary antibody (Cell Signaling Technology, 7074, 7076) diluted in 5% non-fat milk in PBST for 1 hour at room temperature. Membranes were washed three times for 5-10 minutes with PBS at room temperature before being incubated with Pierce ECL (Thermo Fisher Scientific, PI32106) for 2 minutes. Autoradiography films were used to detect the signals.

### Co-immunoprecipitation

Flag-Strep-tagged PPAT and different variants of NUDT5 were over-expressed in PPAT and NUDT5-null HeLa cells as described in **Gene deletion and over-expression**. Cells were scraped in cold PBS on ice and pelleted at 4°C, before being lysed in a buffer containing 20 mM Tris-HCl pH 7.5, 150 mM NaCl, 1 mM EDTA, 1% NP-40, and protease inhibitors (1 µg/ml leupetin, 1 µg.ml pepstatin, 1 mM benazamidine HCl). The cell lysates were subject to three freeze-thaw cycles between liquid nitrogen and a 37 °C water bath, followed by centrifugation at 18,000 x g for 15 min at 4 °C. The supernatants were collected for protein quantification using the DC protein assay kit. After protein normalization, 50 µL supernatants were collected as the input samples mixed with 6x Laemmli buffer and boiled at 95°C for 5 minutes. At least 1 mg protein from the supernatant was incubated with 50 µL 50% Strep-Tactin XT Sepharose chromatography resin (Sigma, GE29401324) on a rotator at 4°C for 2 hours. The samples were spun down at 1,000 x g at 4°C for 30 seconds. The supernatant was aspirated and the resin was washed with 1 mL PBST followed by centrifugation at 1,000 x g at 4°C for 30 seconds. After repeating five times of the wash, 25 µL 2x Laemmli buffer was added followed by 5 minutes of incubation at 95°C to elute the samples three times. Western blots were performed as described in **Immunoblotting.**

### Seahorse XFe96 Respirometry

Oxygen consumption rates were measured using An XFe96 Extracellular Flux Analyzer (Agilent Technologies) as previously described(*8*). In brief, cells were treated with 0.5 µg/mL 6-TG in the morning. 6-8 hours later, cells were re-seeded in a 96-well Seahorse plate with 20,000 cells per well while being exposed to the same concentration of 6-TG for 16-18 hours. Cells were then washed with Seahorse medium three times followed by at least 30 minutes of incubation in a CO_2_-free incubator at 37°C before the analysis. Oxygen consumption rates were normalized to the cell number in each well.

### Immunofluorescence

γH2AX foci were analyzed as previously described(*59*). In brief, cells were seeded on coverslips prior to treatment with vehicle or 0.5 µg/mL 6-TG for 8 or 24 hours. Cells were fixed in 4% paraformaldehyde for 10 minutes followed by 10 minutes of permeabilization in 0.5% Triton X-100 at room temperature. Cells were then blocked with 4% BSA in PBS for 1 hour at room temperature prior to incubation with primary antibodies against γH2AX (1:500, Millipore Sigma) at 4°C overnight. Cells were washed with PBS 3 times before being incubated with secondary antibodies conjugated with fluorophores in the dark for 1 hour and then stained with Hoechst for 5 minutes at room temperature. After 3 x 5 minutes of washing in PBS, the coverslips were mounted and sealed onto glass slides using antifade (Vector Laboratories) and nail polish, respectively. Fluorescent images were captured by Keyence BZ-X700 All-in-one Fluorescence microscope (Keyence). The number of γH2AX and 53BP1 foci were quantified using Image J.

### Purinosome analysis

Live cell fluorescence microscopy analysis of purinosomes was conducted as previously described using transiently expressed FGAMS-GFP (Addgene plasmid # 99107; http://n2t.net/addgene:99107 ; RRID:Addgene_99107) as the purinosome marker(*9, 13*). HeLa cells were cultured in purine-depleted medium (RPMI 1640 medium containing 5% (v/v) dialyzed FBS). Cells were seeded onto 35-mm glass-bottomed dishes (Cellvis) one day prior to transfection. The next day, cells were transfected in OPTI-MEM using FuGENE® 4K reagent (Promega), according to the manufacturer’s guidelines. After incubation with the FuGENE-DNA complexes for 3 hours, cells were washed with PBS and maintained in fresh purine-depleted medium. Imaging was performed 15–20 hours post-transfection at 37°C in a 5% CO_2_ environment using a spinning disk confocal system built on a Leica DMI6000 microscope, equipped with a Yokogawa CSU-X1 spinning disk confocal scanner, a Hamamatsu ImagEMX2 EM-CCD camera, and a Leica 100× oil immersion objective lens (NA = 1.49). 20 µM hypoxanthine was added into the medium to induce purinosome disassembly. Purinosome dynamics were monitored by time-lapse imaging at 30-second intervals for 1 hour. Spinning disk images were processed and analyzed using an in-house Fiji/ImageJ macro script. Briefly, raw images were imported into Fiji/ImageJ, and Labkit(*60*) was utilized to segment purinosomes and generate corresponding masks. Purinosome numbers in each image were quantified using the “Analyze Particles” function of Fiji/ImageJ.

### Statistical analysis

Figures were prepared and statistics were calculated using GraphPad PRISM and R. Statistical calculation details are indicated in the figure legends for each figure. *P < 0.05; **P < 0.01; ***P < 0.001; ****P < 0.0001; NS, not significant (P > 0.05).

---

**Figure S1.**
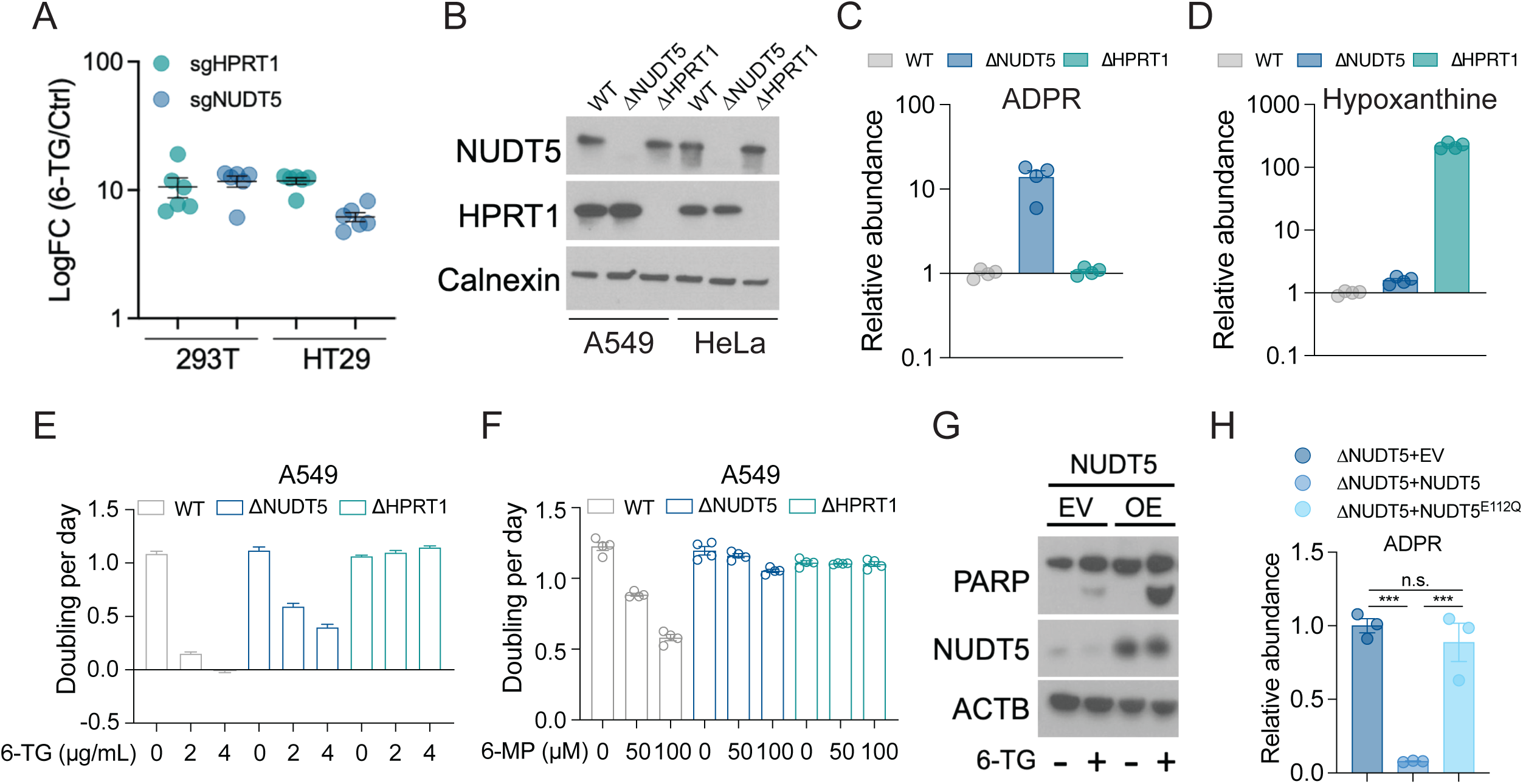
NUDT5 regulates cellular sensitivity to thiopurines independently of its catalytic function. **A.** Enrichment of the indicated gRNA in 6-TG-treated cells during genome-wide CRISPR screen. Data are from Doech et al.(*16*). **B.** Western blot validating depletion of HPRT1 and NUDT5 in HeLa and A549 cells. Calnexin is the loading control. **C-D.** Relative abundance of ADPR (**C**) and hypoxanthine (**D**) in WT, ΔNUDT5, and ΔHPRT1 HeLa cells. Data are means from three replicates. **E-F.** Growth rates of WT, ΔNUDT5, and ΔHPRT1 HeLa cells treated with the indicated doses of 6-TG (**E**) or 6-MP (**F**). Data are from one of three independent experiments. **G.** Western blot assessing cleaved PARP in WT HeLa cells overexpressing an empty vector (EV) or NUDT5. β-actin (ACTB) is the loading control. **H.** Relative abundance of ADPR in ΔNUDT5 HeLa cells that express empty vector (EV), WT NUDT5, or NUDT5^E112Q^. Data are means from three replicates. One-way ANOVA was used for statistical analysis (**H**). ***: P < 0.001; n.s.: P > 0.05. Error bars denote SEM.

**Figure S2.**
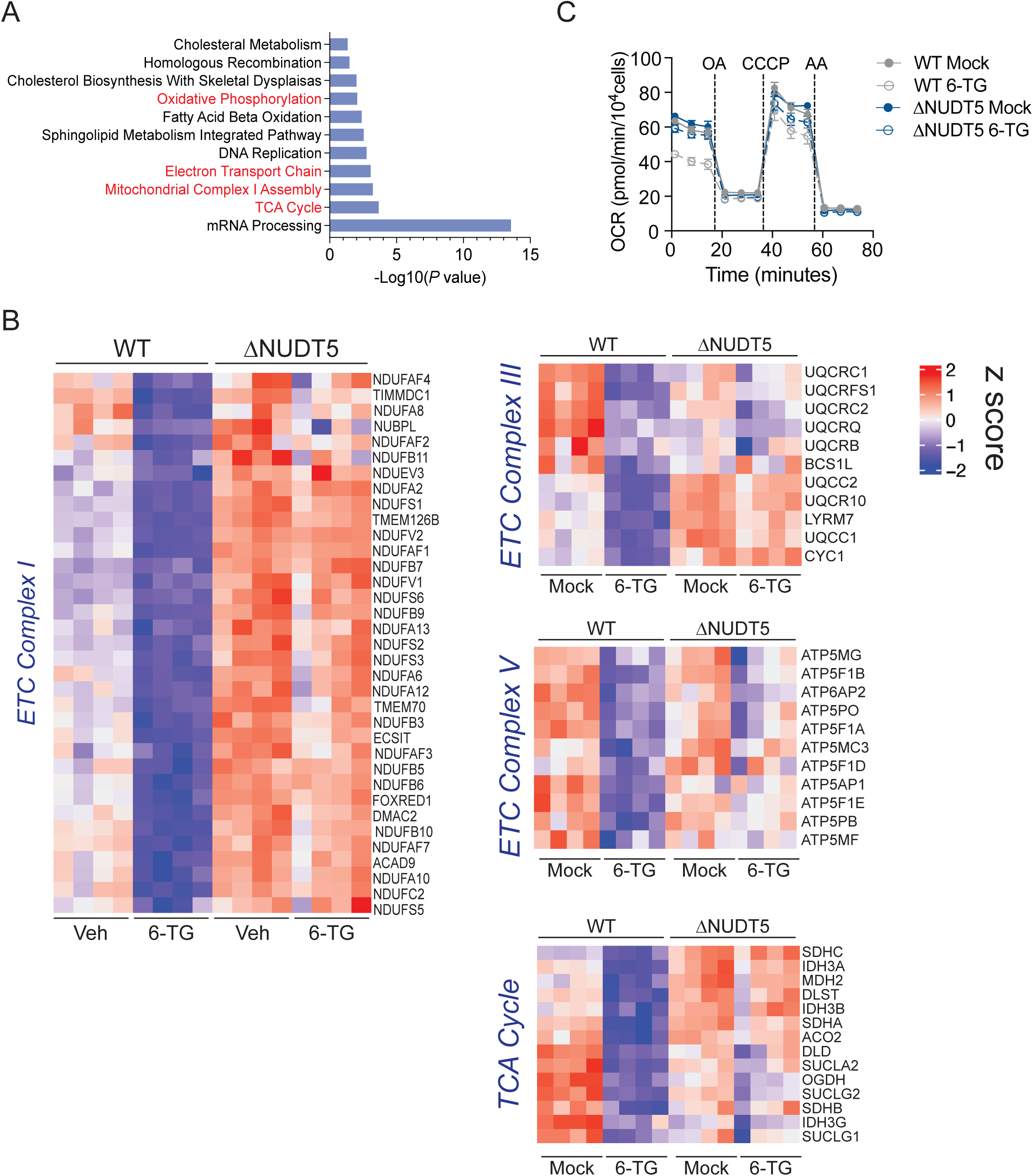
6-TG leads to depleted mitochondrial ETC subunits and reduced respiration in a NUDT5-dependent manner. **A.** Proteomics analysis showing down-regulated pathways in 6-TG-treated WT cells compared to vehicle-treated WT, vehicle-treated ΔNUDT5 and 6-TG-treated ΔNUDT5 cells. **B.** Heatmaps showing abundance of proteins in ETC complexes and the TCA cycle. **C.** Oxygen consumption rates of HeLa cells pre-treated with vehicle or 0.5 µg/mL 6-TG. OA: oligomycin A; CCCP: carbonyl cyanide m-chlorophenylhydrazone; AA: antimycin A. Data are from one of three independent experiments. Error bars denote SEM.

**Figure S3.**
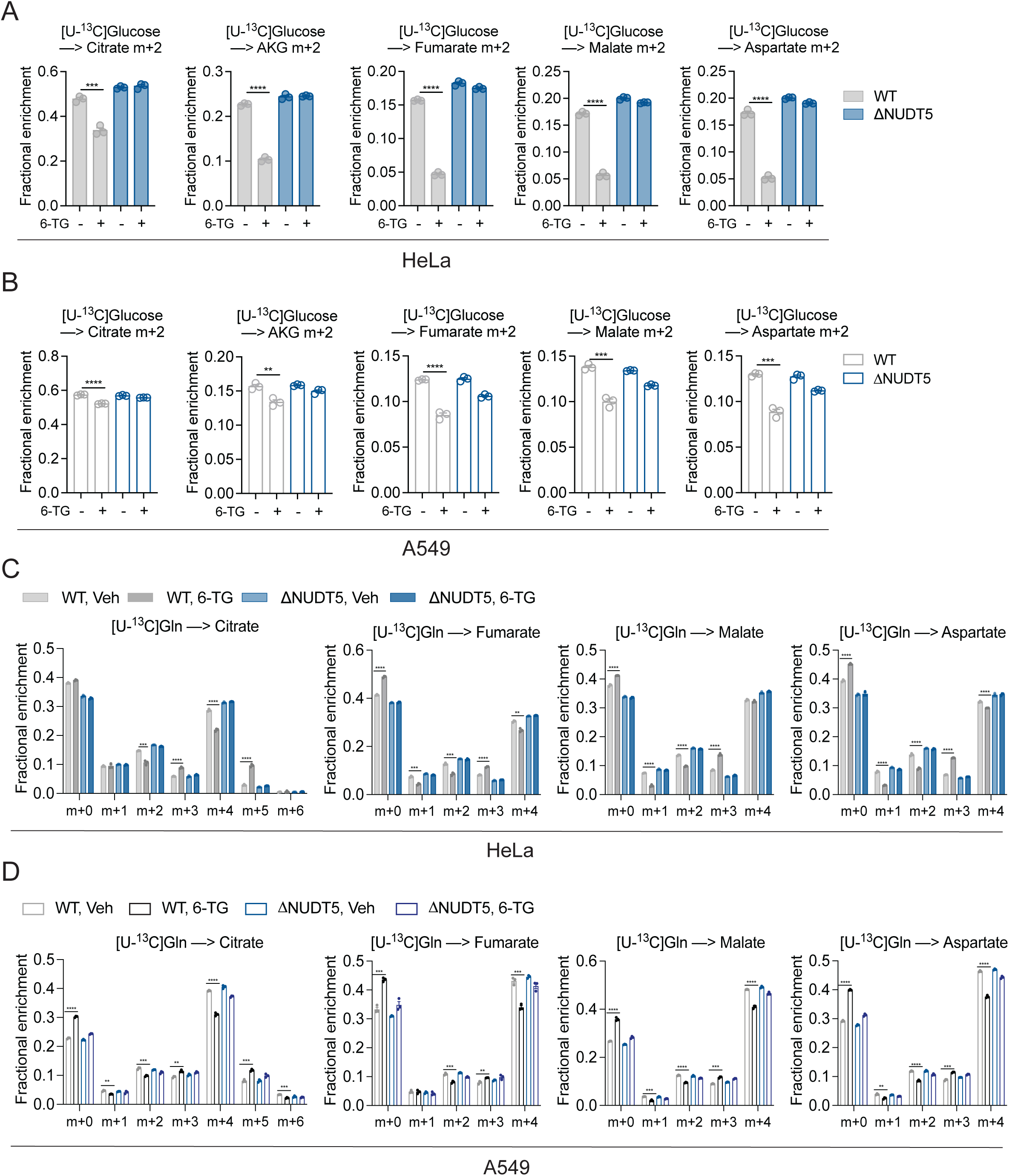
6-TG alters glucose and glutamine metabolism in a NUDT5-dependent fashion. A-B. Isotopologue fractions in citrate, α-ketoglutarate (AKG), fumarate, malate, and aspartate after 6 hours of culture with [U-^13^C]glucose in WT and ΔNUDT5 HeLa (**A**) and A549 (**B**) cells pre-treated with vehicle or 0.5 µg/mL 6-TG for 24 hours. Data are means from three replicates. **C-D.** Isotopologue fractions in citrate, fumarate, malate, and aspartate after 6 hours of culture with [U-^13^C]glutamine in WT and ΔNUDT5 HeLa (**C**) or A549 (**D**) cells pre-treated with vehicle or 0.5 µg/mL 6-TG for 24 hours. Data are means from three replicates. Unpaired (**A-B**) or multiple unpaired (**C-D**) two-sided t tests were used for the statistical analysis. ****: P < 0.0001; ***: P < 0.001; **: P < 0.01. Error bars denote SEM.

**Figure S4.**
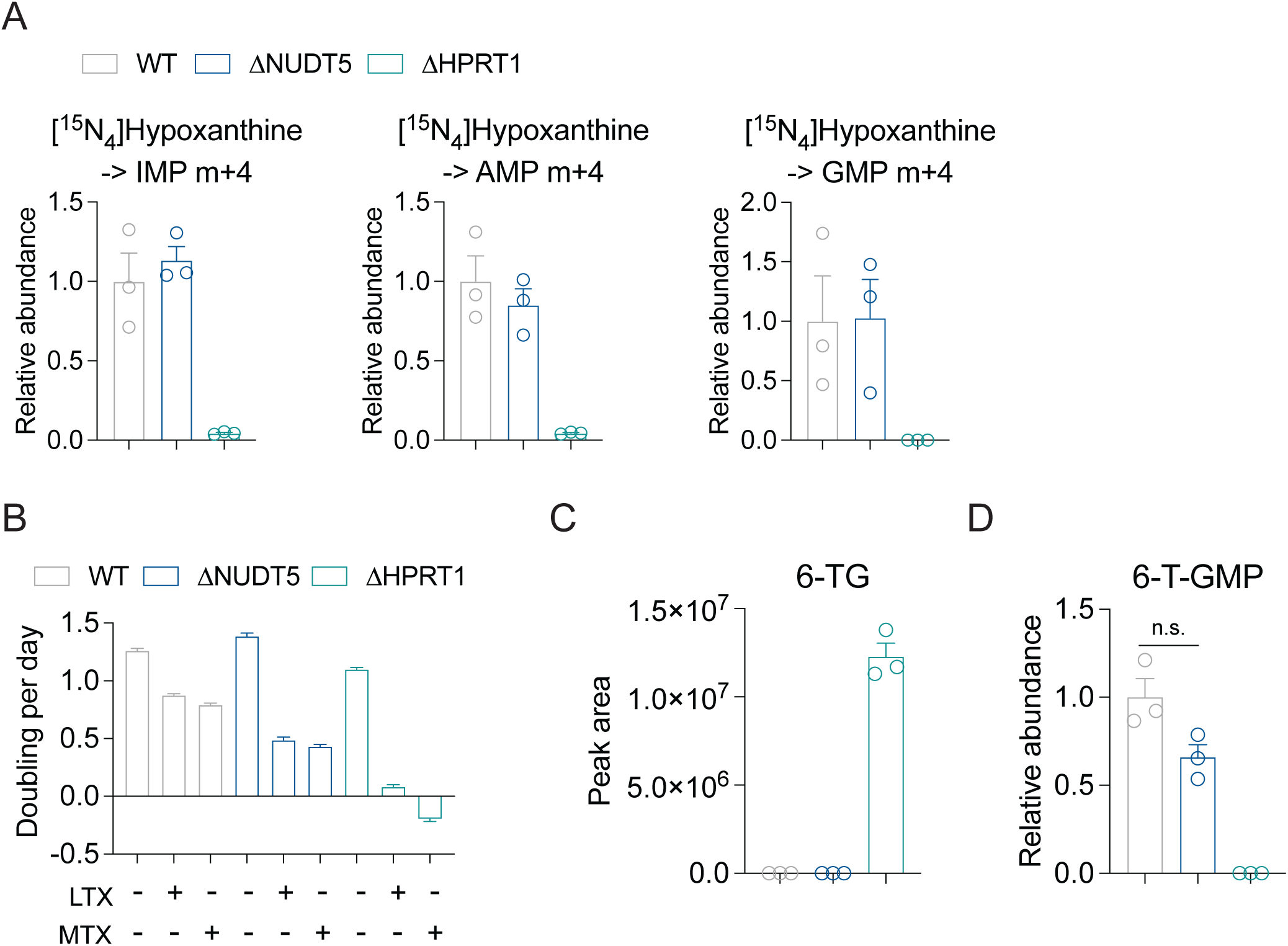
NUDT5 is not required for purine salvage and 6-TG metabolism. **A.** Relative abundance of m+4 IMP, m+4 AMP, and m+4 GMP from [^15^N_4_]hypoxanthine in WT, ΔNUDT5, and ΔHPRT1 A549 cells during 4 hours of tracing. Data are means from three replicates. **B.** Growth rates of WT, ΔNUDT5, and ΔHPRT1 HeLa cells treated with DMSO, 1 µM LTX, or 1 µM MTX. Data are from one of three experiments. **C.** Intracellular 6-TG abundance (peak area) in WT, ΔNUDT5, and ΔHPRT1 A549 cells treated with 0.5 µg/mL 6-TG for 24 hours. No peaks were detected in the WT and ΔNUDT5 cells. Data are means from three replicates. **D.** Relative abundance of 6-T-GMP in WT, ΔNUDT5, and ΔHPRT1 A549 cells treated with 0.5 µg/mL 6-TG for 24 hours. Data are means from three replicates. Error bars denote SEM.

**Figure S5.**
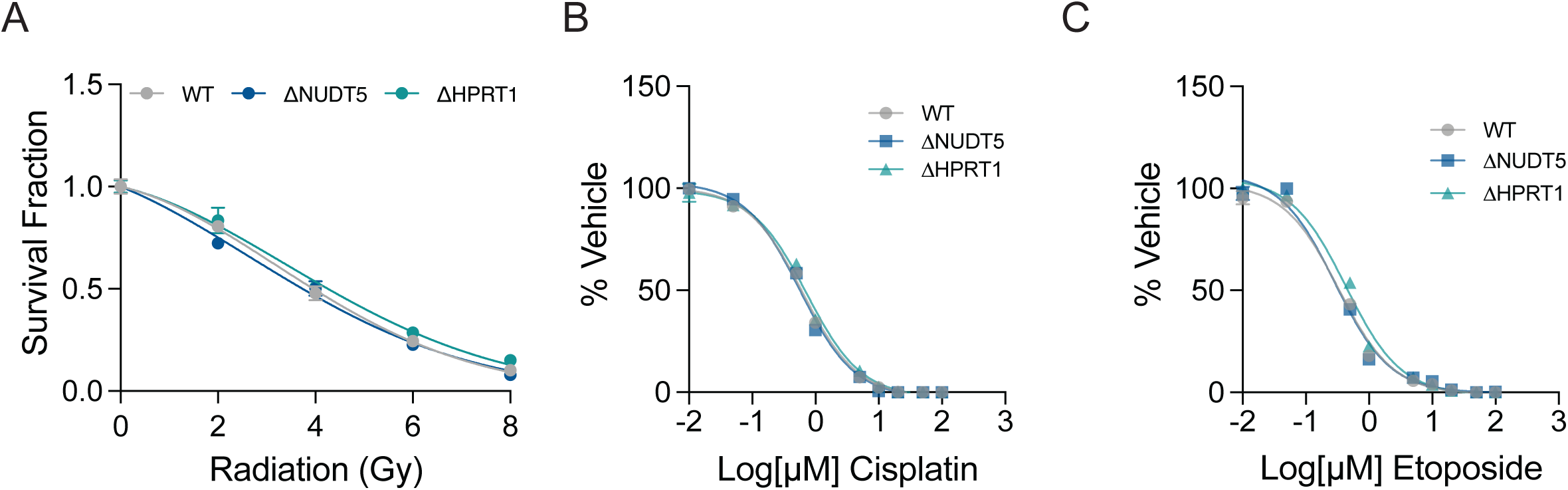
NUDT5 loss does not confer resistance to ionizing radiation or general DNA damaging agents. **A.** Relative survival fraction of WT HeLa cells treated with the indicated doses of ionizing radiation (IR). Data are normalized to the untreated group. Data are means of three replicates. **B-C.** Relative survival of WT HeLa cells treated with the indicated doses of cisplatin (**B**) or etoposide (**C**). Data are normalized to the vehicle-treated groups. Data are from one of three experiments. Error bars denote SEM.

**Figure S6.**
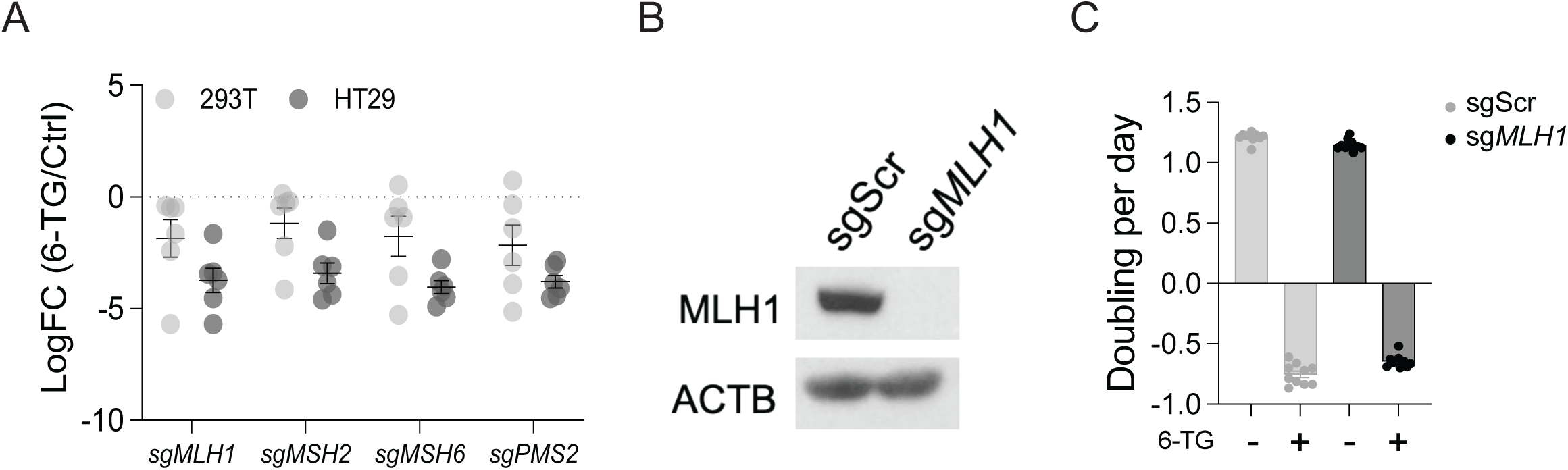
The DNA mismatch repair (MMR) pathway is not involved in NUDT5 depletion-induced thiopurine resistance. **A.** Enrichment of the indicated gRNAs against MMR genes in 6-TG-treated cells during genome-wide CRISPR screen. Data are from Doench et al.(*16*). **B.** Western blot validating depletion of MLH1. β-actin (ACTB) is the loading control. **C.** Growth rates of HeLa cells expressing scrambled gRNA (sgScr) or gRNA against *MLH1* (sg*MLH1*) treated with vehicle or 4 µg/mL 6-TG. Data are from one of three experiments. Error bars denote SEM.

**Figure S7.**
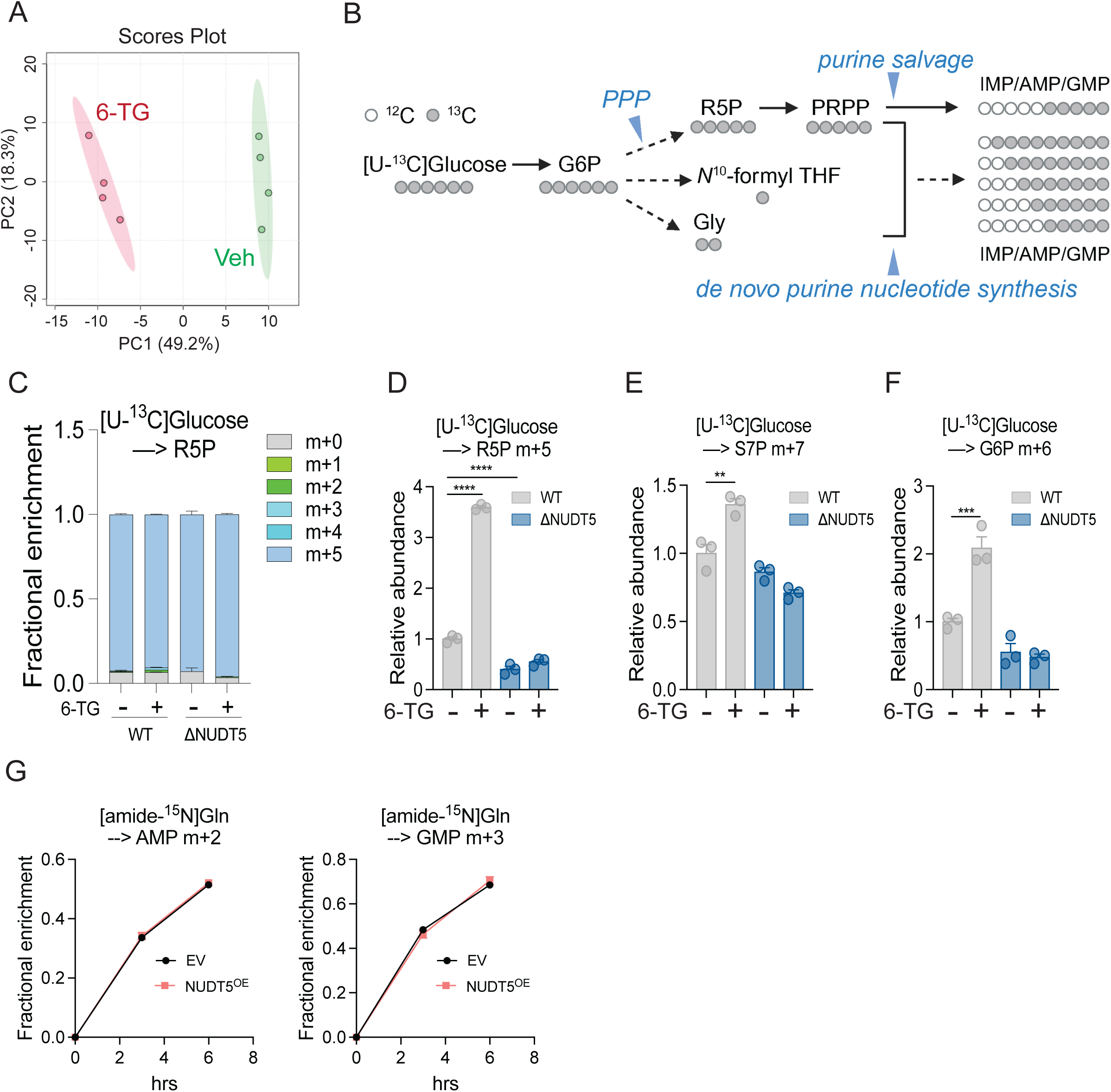
Metabolic effects of 6-TG. **A.** Principal component analysis of metabolomic profiles in WT HeLa cells treated with vehicle (Veh) or 0.5 µg/mL 6-TG for 24 hours. **B.** Schematic illustrating labeling of purine nucleotides from [U-^13^C]glucose. **C-F.** ^13^C labeling in ribose 5-phosphate (R5P) (**C**) and relative abundance of m+5 R5P (**D**), m+7 sedoheptulose 7-phosphate (S7P) (**E**), and m+6 glucose 6-phosphate (G6P) (**F**) from [U-^13^C]glucose during 6 hours of tracing in WT and ΔNUDT5 HeLa cells pre-treated with vehicle or 0.5 µg/mL 6-TG for 24 hours. Data are means from three replicates. **G.** Time-dependent fractional enrichment of m+2 AMP and m+3 GMP from [amide-^15^N]glutamine in WT HeLa cells that overexpress empty vector (EV) or NUDT5. Data are means from three replicates. Unpaired two-sided t tests were used for the statistical analysis (**D-F**). ****: P < 0.0001; ***: P < 0.001; **: P < 0.01. Error bars denote SEM. BioRender was used to generate the illustration in **B**.

**Figure S8.**
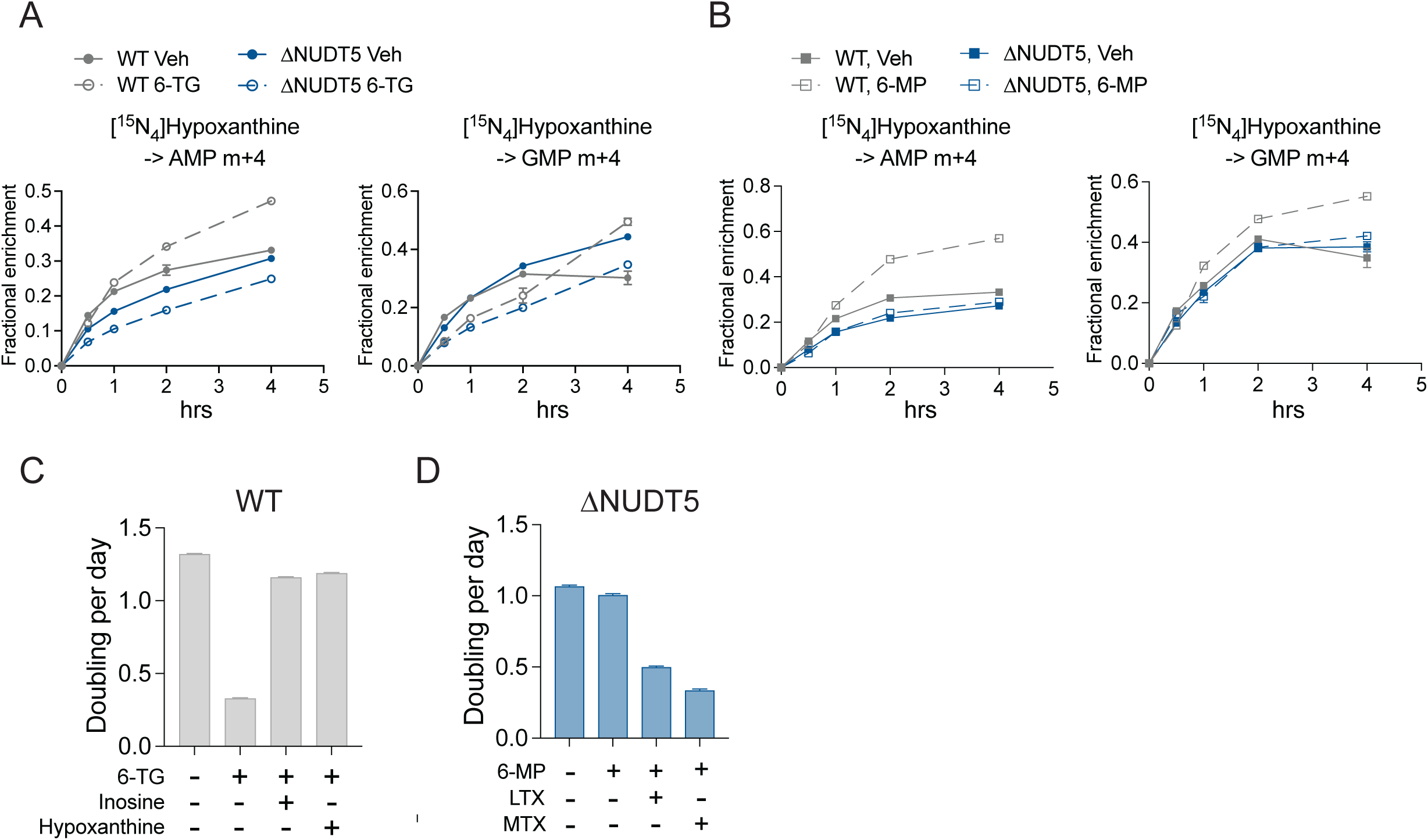
Increased purine salvage in thiopurine-treated WT cells. A-B. Time-dependent fractional enrichment of m+4 AMP and m+4 GMP from [^15^N_4_]hypoxanthine in WT and ΔNUDT5 HeLa cells pre-treated with vehicle, 0.5 µg/mL 6-TG (**A**) or 20 µM hypoxanthine (**B**) for 24 hours. Data are means from three replicates. **C.** Growth rates of WT HeLa cells treated with 0.5 µg/mL 6-TG with or without 50 µM inosine or hypoxanthine. **D.** Growth rates of ΔNUDT5 HeLa cells treated with 0.5 µg/mL 6-TG, 1 µM LTX, 1 µM MTX or the combinations. Error bars denote SEM.

**Figure S9.**
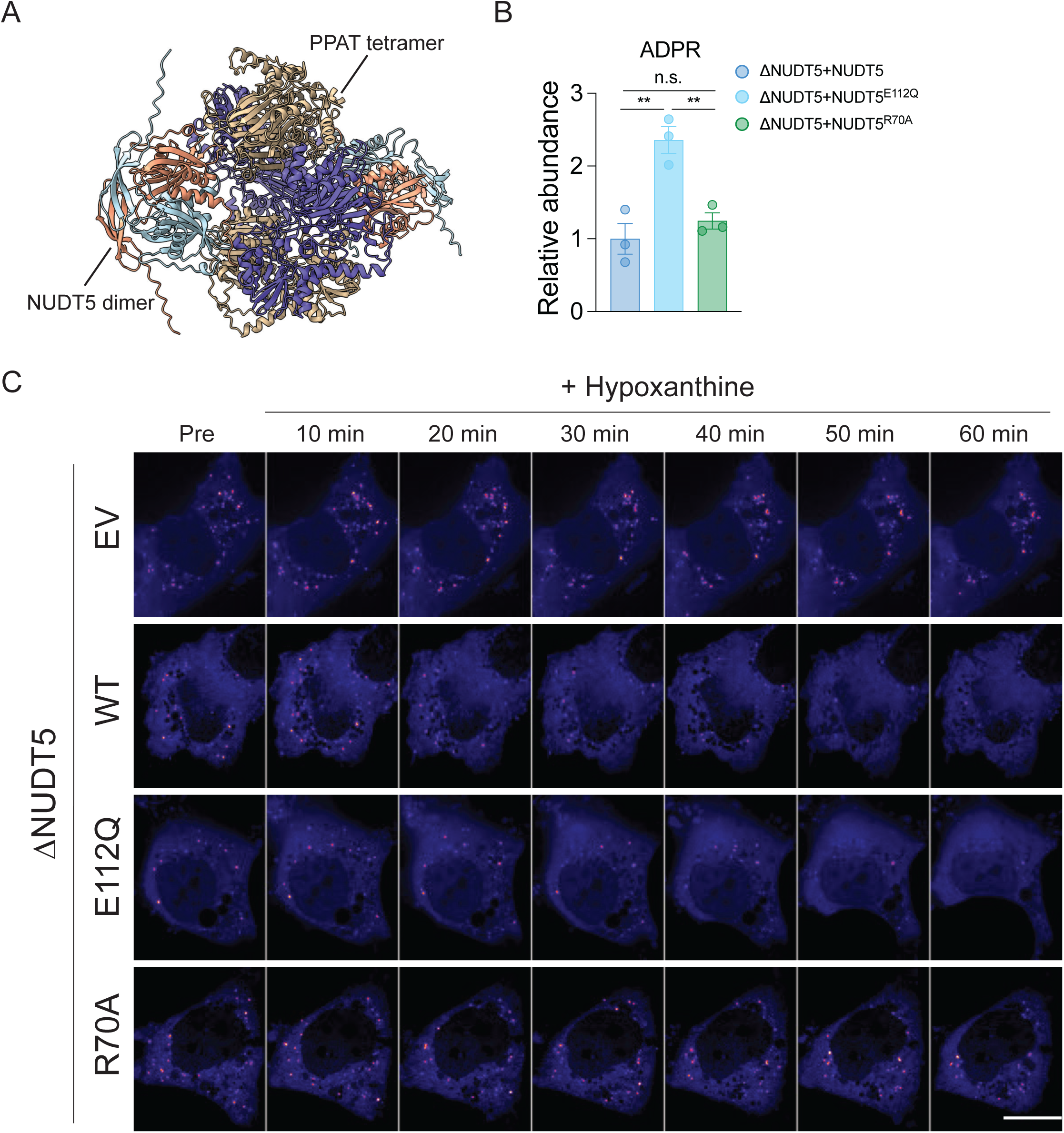
NUDT5 binds PPAT to regulate purinosome dispersal after purine salvage activation. **A.** PPAT tetramer-NUDT5 dimer interaction depicted by the AlphaFold3 structure model. **B.** Relative abundance of ADPR in ΔNUDT5 HeLa cells that express WT NUDT5, NUDT5^E112Q^, or NUDT5^R70A^. Data are means from three replicates. **C.** Time-lapse imaging showing dispersal of purinosomes after treatment with 20 µM hypoxanthine treatment at t=0 minutes in ΔNUDT5 HeLa cells expressing empty vector (EV), WT NUDT5, NUDT5^E112Q^, or NUDT5^R70A^. The puncta indicate GFP-tagged FGAMS. The scale bar represents 10 µm. One-way ANOVA was used for the statistical analysis (**B**). **: P < 0.01; n.s.: P > 0.05. Error bars denote SEM.

**Figure S10.**
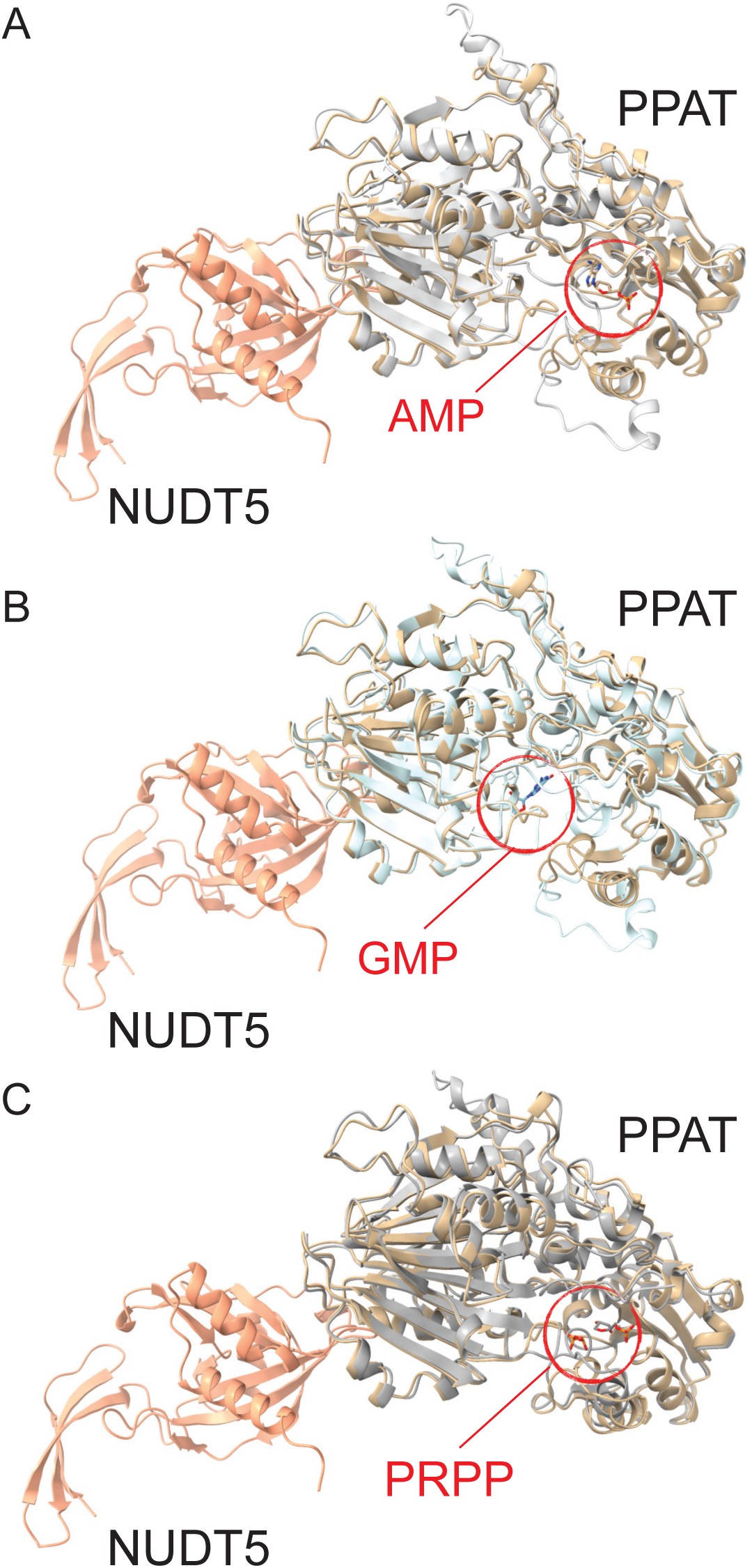
The NUDT5-PPAT interaction is remote from PPAT binding pockets for PRPP and purine nucleotides. A-C. Superposition of the AlphaFold-predicted PPAT structure in its AMP-(**A**), GMP- (**B**), and PRPP- (**C**) bound forms, in complex with NUDT5.

**Figure S11.**
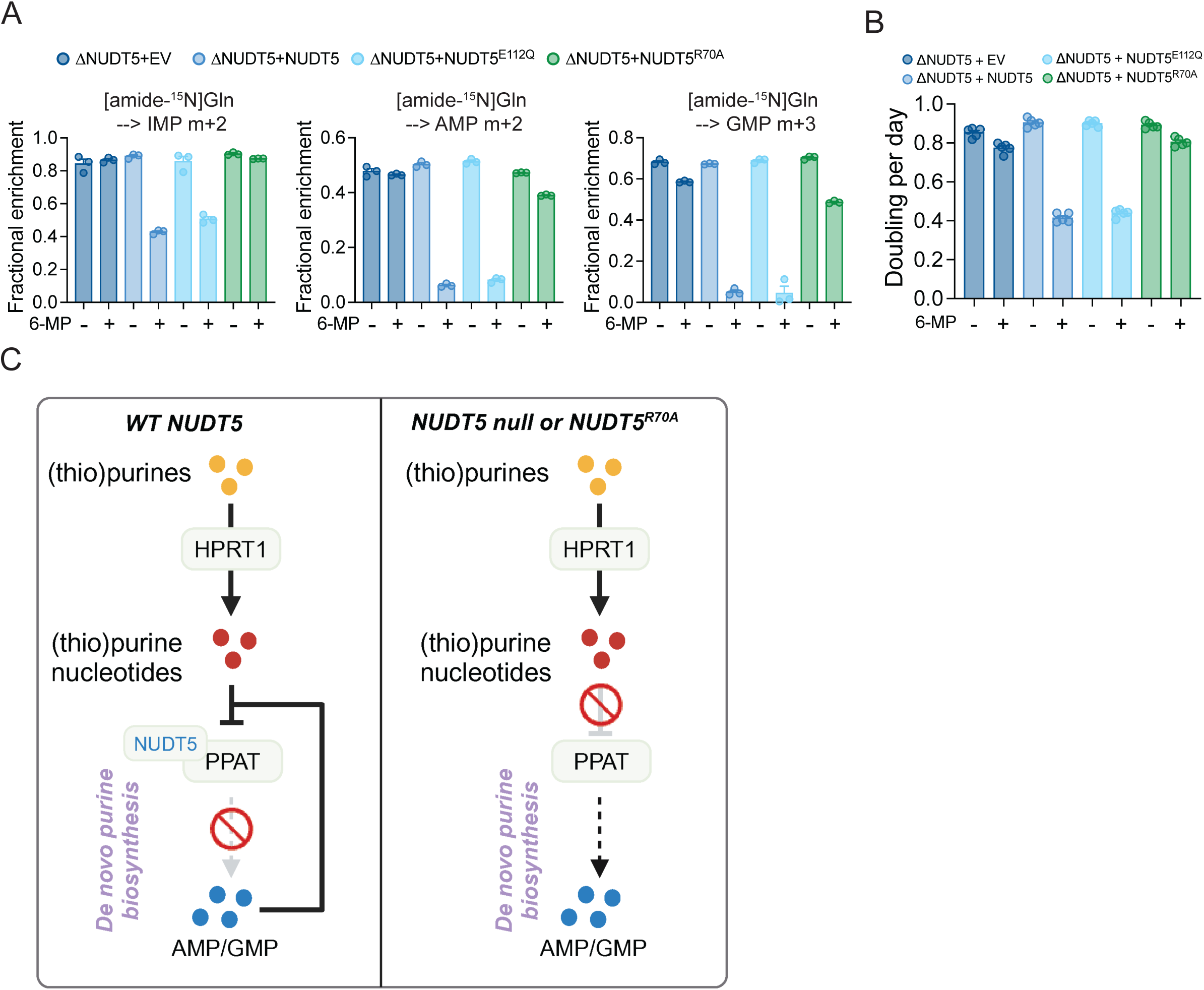
Disruption of the NUDT5-PPAT interaction confers thiopurine resistance. **A.** Fractional enrichment of m+2 IMP, m+2 AMP, and m+3 GMP from [amide-^15^N]glutamine in ΔNUDT5 HeLa cells expressing empty vector (EV), WT NUDT5, NUDT5^E112Q^, or NUDT5^R70A^ during 6 hours of tracing. The cells were pre-treated with 20 µM 6-MP for 24 hours. Data are means from three replicates. **B.** Growth rates of ΔNUDT5 HeLa cells that express empty vector (EV), WT NUDT5, NUDT5^E112Q^, or NUDT5^R70A^ treated with 100 µM 6-MP. Data are from one of three expeirments. **C.** Model of NUDT5 regulation of DNPB. Error bars denote SEM. BioRender was used to generate the illustration in **C**.

